# Beneficial metabolic effects of PAHSAs depend on the gut microbiota in diet-induced obese mice

**DOI:** 10.1101/2023.09.28.558803

**Authors:** Jennifer Lee, Kerry Wellenstein, Ali Rahnavard, Andrew T. Nelson, Marlena M. Holter, Bethany Cummings, Vladimir Yeliseyev, Angela Castoldi, Clary B. Clish, Lynn Bry, Dionicio Siegel, Barbara B. Kahn

## Abstract

Dietary lipids play an essential role in regulating the function of the gut microbiota and gastrointestinal tract, and these luminal interactions contribute to mediating host metabolism. PAHSAs are a class of lipids with anti-diabetic and anti-inflammatory properties, but whether the gut microbiota contributes to their beneficial effects on host metabolism is unknown. Here, we report that treating high fat diet (HFD)-fed germ-free mice with PAHSAs does not improve insulin sensitivity. However, transfer of feces from PAHSA-treated, but not Vehicle-treated, chow-fed mice increases insulin-sensitivity in HFD-fed germ free mice. Thus, the gut microbiota is necessary for and can transmit the insulin-sensitizing effects of PAHSAs in HFD-fed germ-free mice. Functional analyses of the cecal metagenome and lipidome of PAHSA-treated mice identified multiple lipid species that associate with the gut commensal *Bacteroides thetaiotaomicron* (*Bt*) and with insulin sensitivity resulting from PAHSA treatment. *Bt* supplementation in HFD-fed female mice prevented weight gain, reduced adiposity, improved glucose tolerance, fortified the colonic mucus barrier and reduced systemic inflammation versus chow-fed controls, effects that were not observed in HFD-fed male mice. Furthermore, ovariectomy partially reversed the beneficial *Bt* effects on host metabolism, indicating a role for sex hormones in mediating probiotic effects. Altogether, these studies highlight the fact that lipids can modulate the gut microbiota resulting in improvement in host metabolism and that PAHSA-induced changes in the microbiota result in at least some of their insulin-sensitizing effects in female mice.

## INTRODUCTION

Obesity in adults has nearly tripled worldwide in the last fifty years (1), and this is associated with comorbidities and complications that raise long-term healthcare costs and human disease burden (2, 3). The gastrointestinal tract houses the gut microbiota, and its stratified organization of heterogeneous cell types provides both a physical barrier and immune protection to help regulate host metabolism (4–6). Strong evidence supports an essential role for the gut microbiota in the development of obesity (7–12). Obesity caused by high fat diets (HFD) changes gut microbial composition and increases the production of gram-negative bacteria-derived lipopolysaccharide (LPS), which crosses the gut mucosal barrier that has been rendered leaky from the high fat diet. This initiates the chronic low-grade inflammation observed in mice and in many humans with obesity (13–16). Thus, major efforts are focused on developing therapeutic gut-based strategies to restore gut microbiota composition and gut epithelial function to treat metabolic disease. However, the gut microbiota is highly variable in taxonomy and function between individuals, and differences in experimental design further complicate findings and interpretation of results across studies. Furthermore, few studies report sex-specific responses to gut microbial interventions, resulting in major knowledge gaps that limit the identification of optimal treatment strategies for women and men.

Palmitic Acid Hydroxy Stearic Acids (PAHSAs) are a class of lipids with anti-diabetic and anti-inflammatory properties (17–21). PAHSAs were first identified from lipidomics analysis of white adipose tissue from mice overexpressing the glucose transporter, Glut4 (AG4OX) (20). AG4OX mice despite having greater adiposity, have lower fasting glycemia, enhanced glucose tolerance (22, 23), and markedly elevated PAHSA levels in adipose tissue versus controls (20). Circulating and adipose PAHSA levels are low in insulin-resistant people and the levels correlate highly with insulin sensitivity in humans (20). PAHSA treatment regulates multiple components of glucose homeostasis, including augmenting glucose-stimulated insulin secretion through the G-protein coupled receptor GPR40 (19), and regulating insulin action on hepatic glucose production in dietary obese mice (21). PAHSAs also have direct effects in the gastrointestinal tract. Daily PAHSA treatment delayed the onset and reduced the severity of dextran sodium sulfate-induced colitis in mice by modulating innate immune responses and attenuating inflammation (18). Evidence that PAHSAs protect the gut from inflammatory injury along with their beneficial effects on glucose metabolism led us to posit that PAHSAs may alter the gut microbiota in a manner that contributes to their beneficial metabolic effects in dietary obese mice.

In this study, we report that the gut microbiota is essential for and can transmit some of the beneficial metabolic effects of PAHSA treatment in mice. Functional profiling of the cecal metagenome and metabolome from PAHSA-treated mice revealed that specific bacterial species including *Bacteroides thetaiotaomicron (Bt)* and unique signatures of lipid metabolites are altered with PAHSA treatment. Subsequent studies in dietary obese mice demonstrate sex-specific responses to *Bt* supplementation resulting in distinct effects on mucus-producing Goblet cells that line the gut epithelium, intestinal immune phenotypes, and host metabolism. These studies further elucidate the mechanism of action of PAHSAs. They also demonstrate the therapeutic utility of modulating the gut microbiota for the prevention and treatment of obesity and associated metabolic disease and highlight the major role of sex as a biological variable when studying host-microbiome interactions.

## MATERIALS AND METHODS

### Animals

Gnotobiotic (GF) C57BL/6J mice from the Massachusetts Host-Microbiome Center (Brigham and Women’s Hospital, BWH) were used in indicated studies. GF mice at 4-6 weeks of age were co-housed in groups of 3-5 mice per cage in pre-sterilized Optimice cages (for fecal microbiota transplantation (FMT) studies) or in gnotobiotic isolators. C57BL/6J conventional male mice were purchased from Jackson Labs (000664; Bar Harbor, ME, USA) for studies using PAHSA treatment. These mice were maintained in a specific pathogen-free barrier facility with a standard 12:12 light:dark cycle at 22-23°C in the Animal Research Facility at Beth Israel Deaconess Medical Center (BIDMC). These mice were purchased at 4-5 weeks of age and PAHSA treatment was started after one week of acclimatization. Another group of C57BL/6J conventional male mice (Jackson Labs (000664; Bar Harbor, ME, USA)) were fed a HFD to induce insulin resistance, followed by implantation of subcutaneous minipumps to deliver chronic PAHSA treatment (19). In *Bt* supplementation studies, C57BL/6J conventional male and female mice were bred in-house and singly caged in ventilated corn bedding cages with cotton shepherd shacks (BIDMC). *Bt* supplementation was started at 24 weeks of age to determine the effects in chow-fed conventional mice. In a separate cohort, *Bt* supplementation was started at 36 weeks of age to determine the effects in HFD-fed conventional mice. Fecal pellets were collected from GF mice at baseline and serially sampled throughout treatment intervention to ensure gnotobiotic status in experimental mice. GF and conventional mice were fed HFD for 12 weeks prior to starting treatment interventions. Ovariectomy surgeries were performed in mixed genetic background (C57BL/6J and FVB) female conventional mice as described in (24). All experiments were conducted in accordance with protocols for animal use, treatment and euthanasia approved by the Institutional Animal Care and Use Committees of BWH and BIDMC, Boston, MA.

### Rodent diets

Conventional mice housed at BIDMC had *ad libitum* access to drinking water and standard rodent chow diet (Purina Lab Diet 5008). GF mice at BWH had *ad libitum* access to autoclaved drinking water and chow diet (LabDiet 5021, St. Louis, MO, USA). Conventional wild-type and GF mice were fed the same HFD (irradiated, Harlan Teklad Td93075), indicated in the figure legend.

### Tissue Collection

For all GF and conventional mice, trunk blood was collected by decapitation without prior sedation. In conventional mice, tissues were collected in the *ad libitum*-fed state. White adipose tissue (subcutaneous, perigonadal) and brown adipose tissue, gastrointestinal tract (duodenum, jejunum, ileum, colon), cecal contents and fresh fecal pellets (serial collection on indicated days) were snap frozen in liquid nitrogen and stored at −80°C prior to analysis. For studies using GF mice maintained in isolators, feces were collected at baseline, day 7 after the start of chronic treatment and the last day of treatment to confirm germ-free status in mice. The same tissues listed above were collected following a 5-hour food removal. For GF mice in FMT studies maintained in Optimice cages and chronic gavage studies maintained in isolators, feces were collected at days 3 and 7 post-FMT and at terminal tissue collection. The same tissues listed above were collected in the ad-lib fed state (approximately 10AM).

### Body Composition

Body composition for experimental chow and HFD-fed male and female mice was measured by DEXA (dual energy x-ray absorptiometry; Lunar PIXImus, GE Lunar Corp, Madison, WI). Mice were anesthetized with isoflurane. Upon sedation, mice were placed on disposable PIXImus measuring trays for body alignment and fitted with a nose cone administering isoflurane for the duration of the measurement. Mice were returned to home cages for recovery and post-procedure monitoring.

### Serum Biochemistry

Plasma insulin (mouse; Crystal Chem, 90080), total GLP-1 (mouse; Mesoscale Discovery Platform), LBP (Abcam #ab269542), and IL-6 (BioLegend #431307) were measured by ELISA.

### Chronic treatments in experimental mice

*PAHSAs:* 4–5-week-old conventional chow-fed male C57BL/6J mice (Jackson Labs) were singly-housed and treated with either vehicle (50% PEG400: 0.5% Tween-80: 49.5% water) or 5- and 9-PAHSAs (15mg/kg BW of each in vehicle; synthesized by the UC San Diego Center for Drug Discovery Innovation) once daily by oral gavage for 21 days at the ARF at BIDMC. Animals were terminated on day 21 and tissues collected in the 6-hour refed state. Fecal pellets were collected for fecal microbiota transplantation (FMT) studies. Serum and plasma (baseline and terminal time points) and cecal contents were collected for metabolomics and metagenomics analyses. 6–8-week-old GF male and female C57BL/6J mice were co-housed 3-5 mice per cage and maintained in gnotobiotic isolation chambers at BWH. Mice were treated with either vehicle (50% PEG400: 0.5% Tween-80: 49.5% water) or 5- and 9-PAHSAs (dissolved in vehicle; chow dose: 15mg/kg BW of each; HFD dose 45mg/kg BW of each) once daily by oral gavage. Untreated GF control mice did not receive any gavage treatment and maintained their gnotobiotic status.

*Bacteroides thetaiotaomicron VPI-5482 (Bt):* 20-week-old conventional C57BL/6J male and female mice fed chow or HFD were treated with *Bt* inoculum provided by the Harvard Digestive Diseases Center Microbiology Core. Chow mice received 3 treatments by oral gavage per week and HFD-fed mice received daily treatment by oral gavage. Live *Bt* was administered at a dose of 2×10^9^ in 100µL PBS. The heat-killed *Bt* treatment was generated using the same stock of live *Bt* inoculum and heat-treated at 121°C for 15 minutes. All inoculums were prepared under anaerobic conditions and aliquoted into 2mL cryovials for single usage to avoid freeze thawing.

### Fecal Microbiota Transplantation

Fresh fecal pellets from donor conventional vehicle or PAHSA-treated mice were stored at −80°C before being pooled and resuspended with 0.05% cysteine in PBS by Dounce homogenization under anaerobic conditions. Fecal suspensions from donor mice were gavaged in a 200µL volume to 4–6-week-old recipient GF male and female mice maintained in Optimice cages fed chow and HFDs.

### Reagents

The following chemical reagents were purchased from Sigma-Aldrich and used *in vivo:* PEG-400 (81172), Tween-80 (P4780), phosphate buffered saline (PBS; P5368), D-(+) glucose (G8270).

### In vivo Tolerance Tests

Food was removed 5 hours prior to the start of all tolerance tests for all experimental mice. At this time, mice also received their last treatment dose of either vehicle, PAHSAs, or *Bt*. For all tests, baseline glycemia (0 minute timepoint) was measured prior to the administration of glucose (intraperitoneal or oral; 1g/kg BW), insulin (intraperitoneal, dose indicated in figure legend; HumulinR; Lilly, NDC0002-8215-01), or lipid (5µL/g BW). Glycemia with or without blood collection from the tail vein (for insulin secretion assays and serum lipids during a lipid tolerance test) was measured at the indicated time points using a glucometer.

### Gene expression

Fresh ileum and colon from mice were collected and stored at −80°C before being processed for RNA extraction and cDNA synthesis as described (18). Taqman primers for *ppia* (housekeeping gene), *gata3, tbx21, rorc, foxp3, nos2, arg1, il-1β, il-10, il-17, il-23, tnf-α, il-13, il-6, il-4, il-21, ifnγ, rorγt, ahr,* and *occludin* (Applied Biosystems) were amplified for real-time qPCR expression.

### Western blotting

Ileum from experimental mice was processed and protein quantification measured by Bradford assay. 50µg protein was loaded for each tissue sample onto 12% Tris gels (BioRad, Catalog # 567-1044) and transferred onto PVDF membranes. Membranes were incubated with Muc2 (Abcam Catalog # EPR23479-47), IAP (GeneTex Catalog # GTX112100) and GAPDH (Cell Signaling Catalog # 5174s) in 5% BSA-PBS-T at 1:1000 dilution overnight at 4°C followed by 3 washes in PBS-T and 1 hour incubation in secondary antibodies (Fisher; Fluor 488 # PIA32731, Fluor 647 # PIA32733) at room temperature. Blots were imaged on an Odyssey LI-COR imager and quantified by ImageBlot.

### Flow cytometry

Intraepithelial lymphocytes were isolated from mouse colon as described by (25). IELs were stained with the following antibodies: Live/Dead^TM^ Fixable Dead Stain kit (Sigma; L23105), CD45 (V500; BD Biosciences Cat# 561487), CD4 (APC/Cy7; Biolegend Cat# 100414), CD3 (FITC; Biolegend Cat# 100204), CD8a (clone 53-67 PE; Biolegend Cat# 100708), IFNγ (BV421; Biolegend Cat# 505829), TCRγδ (APC; Biolegend Cat# 118116). Where appropriate, IELs were stimulated with Leukocyte Activation Cocktail with BD GolgiPlug^TM^ (BD Pharmingen^TM^, Cat# 550583) for 5 hours prior to cell surface staining, and cells were permeabilized and fixed (Fisher Cat# 50-112-9082; Cat# 50-112-9018, Cat# 005123-43). Unstained, single-stained, and FMO IELs served as controls. Cells were acquired on the Beckman Coulter CytoFLEX LX followed by data analysis on FlowJo v10.8.1.

### Short-chain fatty acid measurements

Cecal samples were kept frozen at −80°C until analysis. HPLC grade water was added to each sample at a volume equal to the weight of the cecal content. Samples were vortexed for 5 minutes until the material was homogenized and then centrifuged at 10,000g for 5 minutes. 100µl of the clear supernatant was transferred into a new Eppendorf tube for further processing. The pH of the suspension was adjusted to 2-3 by adding 10µL of 50% sulfuric acid, 10µL of the internal standard (1% 2-methyl pentanoic acid solution) and 100µL of ethyl ether anhydrous were added to each sample. The tubes were vortexed for 2 minutes and then centrifuged at 5000g for 2 minutes. The upper ether layer was transferred into the Agilent sampling vial for analysis. Standard solutions comprised of a volatile acid mix containing 10 mM of acetic, propionic, isobutyric, butyric, isovaleric, valeric, isocaproic, caproic, and heptanoic acids was used (Supelco CRM46975, Bellefonte, PA). A standard stock solution containing 1% 2-methyl pentanoic acid (Sigma-Aldrich St. Louis, MO) was prepared as an internal standard control for the extractions. Chromatographic analyses were carried out using an Agilent 7890B system with flame ionization detector (FID) (Agilent Technologies, Santa Clara, CA). High-resolution gas chromatography capillary column coated with 0.25µm film thickness was used (DB-FFAP) for the detection of volatile acids (Agilent Technologies, Santa Clara, CA). The oven temperature was 145°C and the FID and injection port were set to 225°C. Nitrogen was used as the carrier gas. 5µl of extracted sample was injected for analysis and the run time was 11 minutes. Chromatograms and data integration were carried out using the OpenLab ChemStation software (Agilent Technologies, Santa Clara, CA). SCFAs were quantified using a 100µL of the standard mix and processed as described for the samples. The retention times and peak heights of the acids in the standard mix were used as references for the sample unknowns. These acids were identified by their specific retention times and the concentrations determined and expressed as mM concentrations per gram of fecal material.

### Metabolomics

Lipids and nonpolar metabolites were extracted from 10µL plasma or cecal homogenate using 190µL of isopropanol containing an internal standard, separated using reversed phase C8 ultrahigh performance chromatography (U-HPLC), and analyzed using high resolution, full scan mass spectrometry in the positive ion mode. Measurements were performed by the Broad Metabolomics Platform. Raw LC-MS data were acquired to the data acquisition computer interfaced to each LC-MS system. Targeted data processing of known metabolites was determined using TraceFinder (v 3.3, Thermo Scientific) software and compound identities are confirmed using reference standards and reference samples included in each analysis queue. Concatenated data from each of the methods is expressed as individual samples in columns and metabolite abundances in rows.

### Microbiome Sequencing

Cecal contents from conventional C57BL/6J mice (Jackson labs) treated with vehicle or PAHSAs were processed after 21 days of treatment for metagenome sequencing (MRDNA, Texas, USA). Fecal pellets from *Bt* supplementation studies were collected after 61 days of treatment and processed for metagenome sequencing (The Genomics Core, George Washington University).

### Microbiome taxonomic and functional profiling

To ensure quality, raw reads from the whole metagenomic shotgun sequencing of these samples were filtered using the Kneaddata quality control software (26). This produced trimmed reads devoid of the host reads. The cleaned reads were then categorized for relative abundance using MetaPhlAn embedded in HUMAnN (27), where they were mapped against reference genomes for microbial species and then functional profiling of these reads was profiled by multi-omics correlation analysis.

### Multi-omics Correlation Analysis

Correlations between microbial species abundance and metabolite intensities were assessed using the *’btest’* Python package (28). ‘btest’ is a method to link, rank, and visualize associations among omics features across multi-omics datasets. btest conducts correlation tests for omics features across two paired omics datasets that share common samples. These paired datasets contain different omics measurements but originate from the same set of samples. Within *’btest’*, the p-value and correlation are determined using the Spearman correlation test. Adjustments for multiple testing are made using the Benjamini-Hochberg procedure, aiming for a significant association at a False Discovery Rate threshold of 0.05.

### Histology and Imaging

Colon sections were fixed in Carnoy’s solution for 24 hours followed by serial dehydration and paraffin-embedded for cross-sectional tissue processing for AB/PAS staining. 5-8 images were taken for each colon and the average mucosal thickness from 10-20 cross-sectional images measured by Scion Image is reported for each experimental mouse. Ileum sections were fixed in 10% formalin and cross-sectioned for AB/PAS staining. Positively stained goblet cells were counted in 20-50 villi per ileal section and the average number of goblet cells was reported for each experimental mouse. All images were taken by light microscopy (Zeiss AxioImager M1 Epifluorescence and Brightfield Microscope) at 10X and 20X magnification where indicated.

### Quantification and Statistical Analysis

All data are presented as means±SEM. Significance was assessed by either Student’s t-test, one- or two-way ANOVA with Tukey post-hoc test for multiple comparisons where appropriate except for the btest as explained above. Bar blots and line graphs were generated with GraphPad Prism and differences were considered significant when p<0.05. GraphPad Prism was used to generate scatterplots and to determine correlation coefficients and p-values using the Spearman correlation test. Statistical parameters can be found in the figure legends. Stacked bar plots representing gut microbiota abundance levels were generated by Microsoft Office Excel. Heatmaps and PCA plots related to gut microbiota abundances and species variance analysis were generated by Qlucore Omics Explorer 3.7(24) and differences were considered significant when p<0.05 followed by post-hoc adjustment by variance with q<0.05.

## RESULTS

### PAHSA treatment in mice increases insulin sensitivity and alters the gut microbiota and plasma and cecal metabolomes

The effects of PAHSA treatment on gut microbiota function and community composition have not been reported. To determine this, we measured the cecal and plasma metabolomes and cecal microbiome in conventional chow-fed mice (Figure 1A). Consistent with previous reports, PAHSA treatment had no effect on body weight (19, 21) (Figure 1B) but improved insulin sensitivity as early as 13 days of treatment (Figure 1C). PAHSAs had major effects on lipid metabolite levels in plasma (Figure 1D). PAHSA-treated mice had increased circulating levels of seven cholesterol ester (CE) species (16:0, C18:1, C18:2, C18:3, C20:3, C20:5, C22:6) and low levels of eighty plasma lipid metabolites compared to controls, with 65 of these species being triacylglycerols (TAGs), ten DAGs (diacylglycerides; C34:3, C36:2, C36:3, C36:4, C38:4, NH4-C34:3, NH4-C36:2, NH4-C36:3, NH4-C36:4, NH4-C38:4), three Pes (phosphatidylethanolamine; C36:1, C36:2, C36:3), and two PCs (phosphatidylcholine; C34:0, C36:4 PC-A) (Figure 1D). By contrast, only thirty-eight lipid-related cecal metabolites were differentially regulated in PAHSA-treated versus control mice (Figure 1E). Of these cecal lipid metabolites, the majority increased with PAHSA treatment, including five PCs (C34:1, C36:1, C36:2, C38:4, C40:6), six PEs (C36:1, C36:2, C36:4, C38:2, C38:4, C38:6), sixteen TAGs (between C48-C56), and three plasmalogens (C34:2 PE, C38:7 PC, C40:7 PE) (Figure 1E).

**Figure 1.**
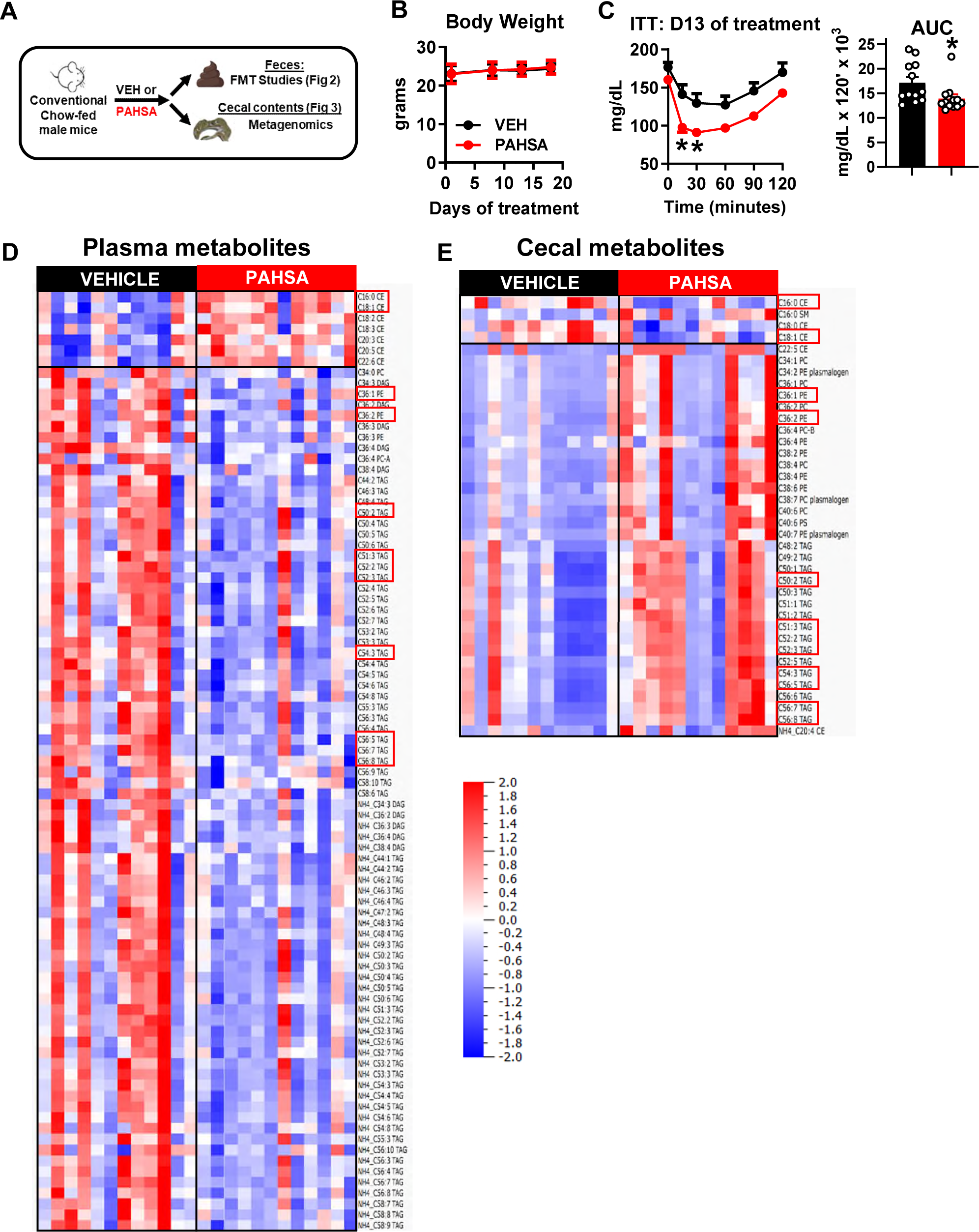
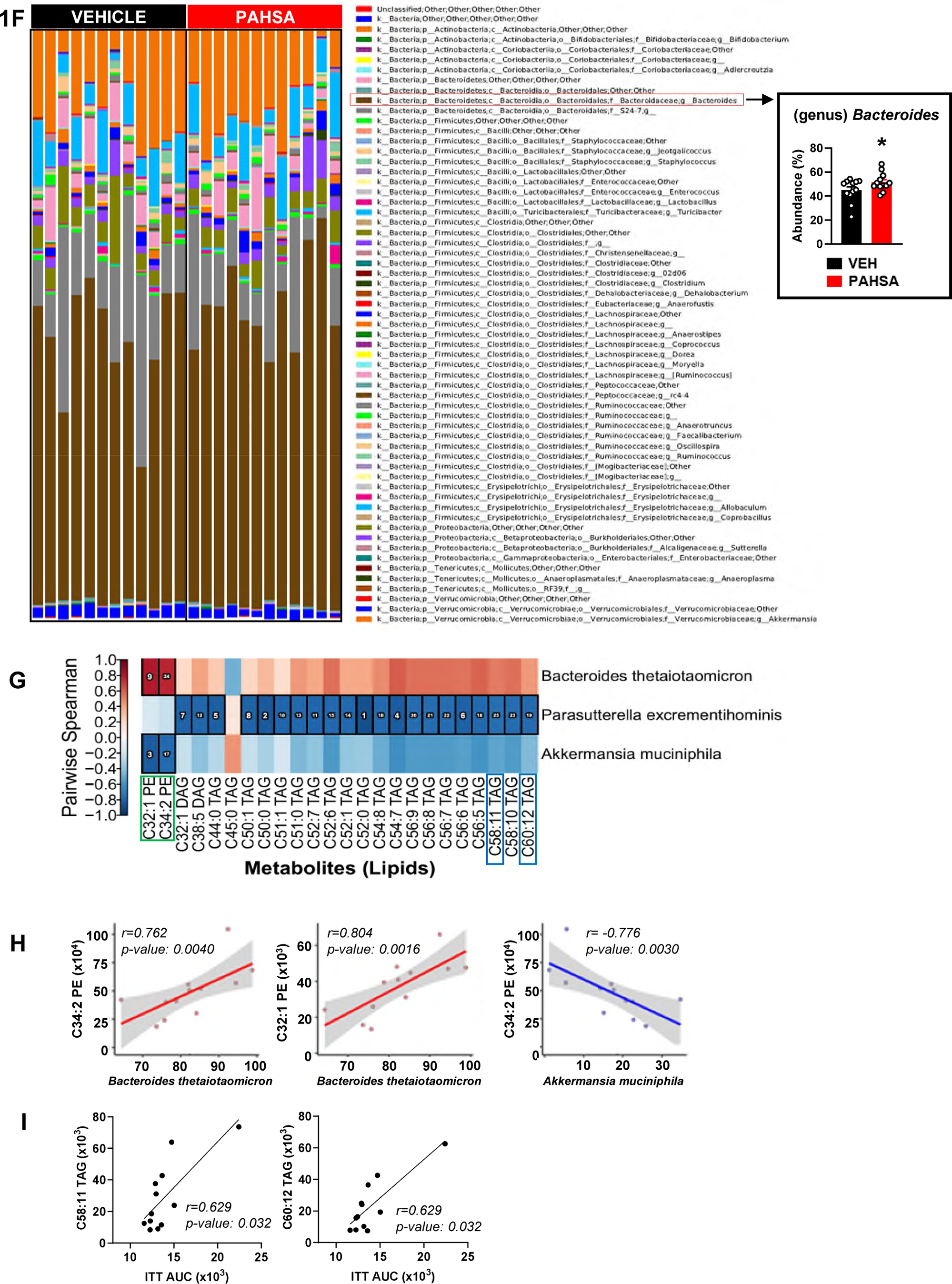
PAHSA treatment in mice increases insulin sensitivity and alters the gut microbiota and plasma and cecal metabolomes. **(A)** Experimental design in which conventional chow-fed male mice were treated once daily with oral vehicle or PAHSAs. Effects of treatment on **(B)** body weight, **(C)** insulin sensitivity after 13 days of treatment, with area under the curve (AUC) (*p<0.05 PAHSA vs VEH). Heatmaps of abundance levels of **(D)** plasma metabolites, and **(E)** cecal metabolites. Each column represents one mouse. Stacked bar graph of abundance levels of gut microbiota at the genus level in vehicle and PAHSA-treated mice (**F**). Bar graph of abundance level of **(F)** *Bacteroides* at the genus level, (*p<0.05 PAHSA vs VEH). **(G)** Heatmap showing that functional analysis using *’btest’* Python package which links, ranks, and visualizes associations among omics features across multi-omics datasets, identifies *Bacteroides thetaiotaomicron* (*Bt)* to most strongly associate with a subset of cecal lipid metabolites from PAHSA-treated mice. **(H)** Correlation graphs show associations of *Bt* with C34:2 PE and C32:1 PE, and *Am* with C34:2 PE. **(I)** Correlation graphs show associations of ITT AUC with C58:11 TAG and C60:12 TAG. n=12/group. Statistical analyses for 1C and 1F were conducted using repeated-measures two-way ANOVA followed by Tukey’s post-hoc test and Student’s t-test. btest was used in 1H using data in 1E and the bar graph in 1F. The Spearman correlation test was used in 1I using the bar graph in 1C and using the data in 1E. The Spearman correlation test was used in 1H and 1I to calculate r- and p-values. Data in 1B, 1C, and the bar graph in 1F are presented as means ± SEM. Red boxes in 1D and 1E indicate metabolites that are changed in both plasma and cecal contents, although sometimes in the opposite direction, from the same experimental mice. In 1G, green boxes indicate metabolites that positively correlate with *Bt* abundance levels, and blue boxes indicate metabolites that positively correlate with ITT AUC of PAHSA-treated mice.

Twelve of these metabolites were changed in both plasma and cecum of PAHSA-treated mice, but not always in the same direction. These included two CEs (C16:0, C18:1), two PEs (C36:1, C36:2), and eight TAGs (C50:2, C51:3, C52:2, C52:3, C54:3, C56:5, C56:7, C56:8). For example, levels of C16:0 and C18:1 CEs were higher in plasma and lower in cecum of PAHSA-treated mice, and PEs and TAGs were lower in plasma and increased in cecum of PAHSA-treated mice (Figures 1D and 1E). These data indicate that a sub-set of lipid metabolites are differentially regulated between the cecum and circulation.

16S rRNA gene phylotyping analyses of cecal contents showed comparable abundance levels among operational taxonomic units (OTU) for most genera between vehicle and PAHSA-treated mice, with the exception of *Bacteroides* OTU, which increased with PAHSA treatment (Figure 1F). We applied the HUMAnN2 analytical method (27) to comprehensively integrate species-level taxonomic information with the cecal metabolome to gain a better understanding of gut microbiome function in insulin-sensitive PAHSA-treated mice. Twenty-four lipid metabolites (two PEs (C32:1 and C34:2), two DAGs (C32:1 and C38:5), and twenty TAGs (>C44:0 and <C60)) strongly associated with three microbial species in PAHSA-treated mice: *Bacteroides thetaiotaomicron (Bt* - positive*), Parasutterella excrementihomonis (Pe –* negative), and *Akkermansia muciniphila (Am –* negative) (Figure 1G heatmap). Correlation dot plots show that the two PE species strongly correlated with *Bt* but not to *Am* (Figure 1H). Furthermore, within the subset of twenty-four lipid metabolites (Figure 1G), two lipid metabolites (C58:11 TAG, C60:12 TAG) positively correlated with the ITT AUC of insulin sensitive PAHSA-treated mice (Figure 1I). These data demonstrate that PAHSAs have specific effects on the gut microbiota and lipid metabolites, and that these gut microbe-metabolite signatures may contribute to the improved insulin sensitivity observed in PAHSA-treated mice.

In the same cohort of mice, PAHSA treatment increased ileal expression of *gcg* and the Goblet cell marker *clca1* (Figure S1A). In another cohort of mice fed a HFD, PAHSA treatment reduced the expression of pro-inflammatory cytokines and increased the expression of *occludin*, a tight junction marker in ileum (Figure S1B). Altogether, these data indicate that PAHSAs have direct beneficial effects on the gut epithelium. PAHSA treatment did not alter levels of gut microbe-derived short-chain fatty acids (SCFA) in cecum (data not shown). By contrast, in another cohort of chow-fed mice that were refed for 4 hours, PAHSA treatment for 28 days raised fecal propionate levels (data not shown). These differences in SCFA levels may be due to the site in the gastrointestinal tract from which the sample was taken, and the fed versus fasted/refed state of the animal.

### The beneficial PAHSA effects on glucose homeostasis are transmissible by fecal microbiota transplantation

Next, we aimed to determine whether the PAHSA effects to improve host metabolism can be transmitted by fecal microbiota transplantation (FMT). Donor feces from conventional male mice treated with vehicle or PAHSAs (Figure 1A) were used for fecal microbiota transplantation (FMT) into recipient GF-HFD male mice (Figure 2A). Similar to the fact that PAHSA treatment did not affect body weight in donor mice, there was no effect of PAHSA-FMT on body weight in recipient mice compared to Vehicle-FMT controls (Figures 2B-C). However, PAHSA-FMT recipient male mice were more glucose tolerant and insulin sensitive than Vehicle-FMT recipient mice, with responses comparable to GF control mice receiving no FMT (Figures 2D-F). By contrast, there was no effect of PAHSA-FMT on any of these metabolic outcomes in GF-HFD female mice (Figures S2A-F). This indicates sex-specific responses in HFD-fed GF mice to FMT from PAHSA-treated mice. These data demonstrate that FMT from PAHSA-treated chow-fed mice can transmit the PAHSA-mediated improvements in glucose metabolism to male GF-HFD mice.

**Figure 2.**
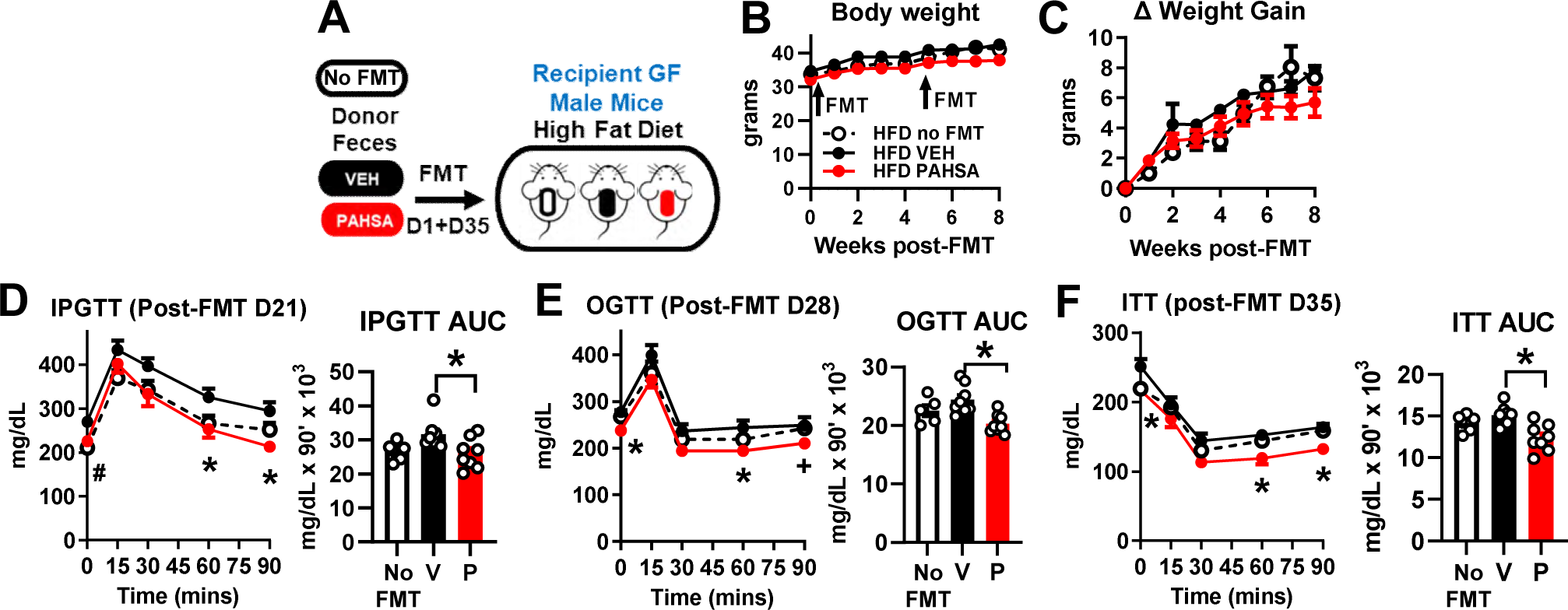
The beneficial PAHSA effects on glucose homeostasis are transmissible by fecal microbiota transplantation. **(A)** Experimental design for fecal microbiota transplantation (FMT) studies in which fecal contents from donor chow-fed male mice treated with vehicle (V) or PAHSAs (P) were orally administered to recipient germ-free (GF) HFD-fed male mice. GF-HFD mice that did not receive FMT (HFD no FMT) served as controls. Recipient GF-HFD mice received two oral doses of fecal inoculum on days 1 and 35. Effects of FMT in recipient GF-HFD male mice on **(B)** body weight, **(C)** weight gain, **(D)** intraperitoneal glucose tolerance 21 days post-FMT, **(E)** oral glucose tolerance 28 days post-FMT, and **(F)** insulin sensitivity 35 days post-FMT. *p<0.05 GF-HFD mice receiving FMT from VEH-treated chow-fed mice vs GF-HFD mice receiving FMT from PAHSA-treated chow-fed mice, #p<0.05 GF-HFD mice receiving FMT from VEH-treated chow-fed mice vs all groups. +p<0.05 GF-HFD mice receiving FMT from PAHSA-treated mice vs GF-HFD mice receiving no FMT. n=5 GF-HFD receiving no FMT, n=8 GF-HFD receiving FMT from VEH-treated mice, n=8 GF-HFD mice receiving FMT from PAHSA-treated mice. All data presented as means ± SEM. Statistical analysis conducted using one- or two-way ANOVA with repeated measures followed by Tukey’s post-hoc test where appropriate.

### The gut microbiota is necessary for PAHSA effects to improve glucose homeostasis in HFD-fed mice

To determine whether the gut microbiota contributes to the anti-diabetic effects of PAHSAs, we treated germ-free (GF) HFD-fed female and male mice with oral PAHSAs or vehicle for four weeks (Figure 3A, S3A). Consistent with previous studies in conventional dietary obese mice (19, 21), there was no effect of PAHSA treatment on body weight in GF-HFD female mice (Figure 3B-C). In addition, PAHSA treatment did not improve glucose tolerance (Figure 3D) or insulin sensitivity (Figure 3E) in GF-HFD female mice. These effects were also observed in GF-HFD male mice (Figures 3F-J). These data indicate that PAHSA effects to improve glucose homeostasis in HFD-fed mice requires the gut microbiota.

**Figure 3.**
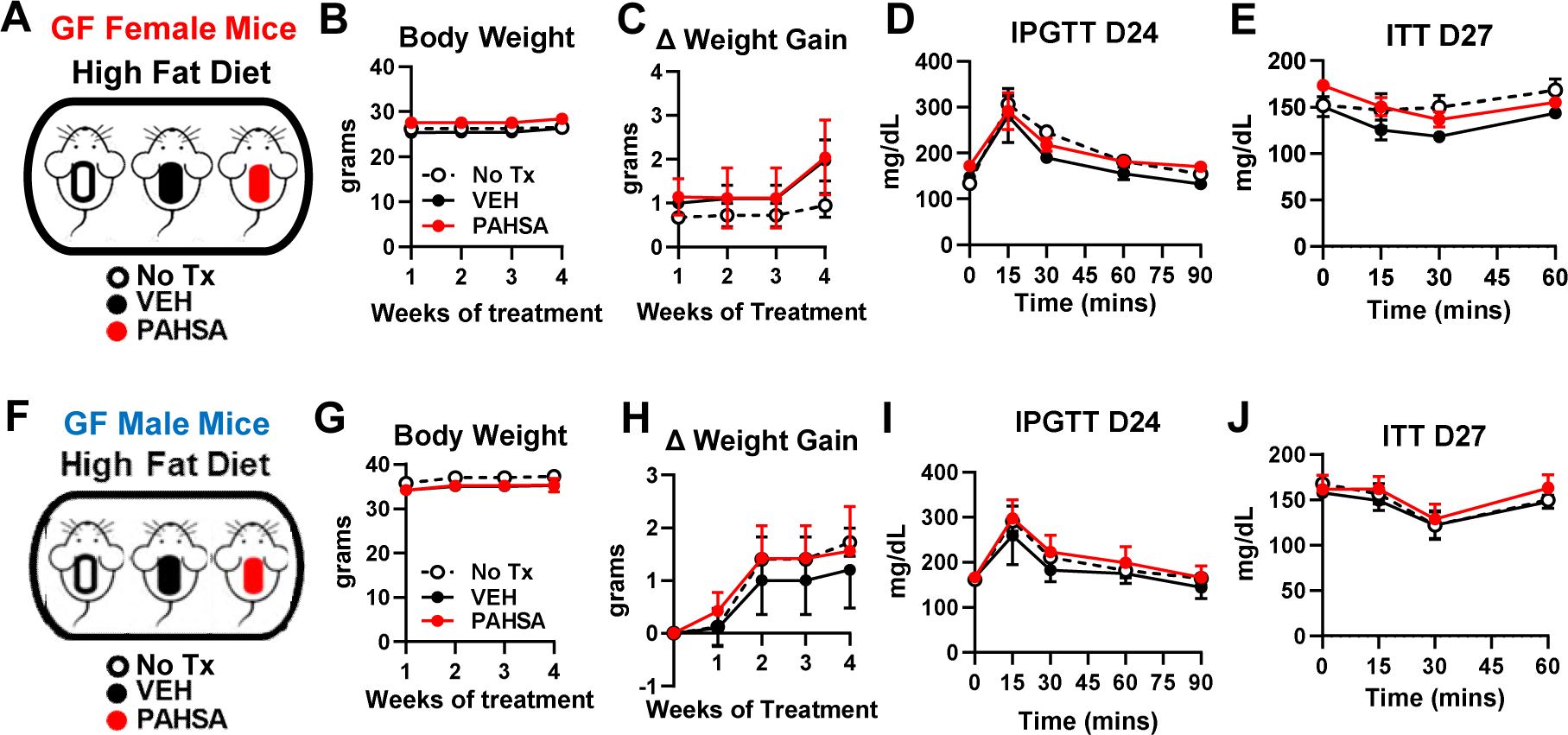
The gut microbiota is necessary for PAHSAs to improve glucose homeostasis in HFD-fed mice. **(A)** Experimental design for HFD-fed GF female mice receiving no treatment (No Tx) or treated with oral vehicle (VEH) or PAHSAs. Effects of once daily oral vehicle or PAHSA treatment in GF-HFD female mice on **(B)** body weight, **(C)** weight gain, **(D)** intraperitoneal glucose tolerance after 24 days of treatment, and **(E)** insulin sensitivity after 27 days of treatment. **(F)** Experimental design for GF-HFD-fed male mice receiving no treatment (No tx) or treated once daily with oral vehicle (VEH) or PAHSAs. Effects of once daily oral vehicle or PAHSA treatment in male GF-HFD mice on **(G)** body weight, **(H)** weight gain, **(I)** intraperitoneal glucose tolerance after 24 days of treatment, and **(J)** insulin sensitivity after 27 days of treatment. n=4-5/group in female mice and n=3-5/group in male mice. All data presented as means ± SEM.

### The gut microbiome has sex-specific responses to *Bacteroides thetaiotaomicron* supplementation in mice

Functional profiling of the gut microbiota from PAHSA-treated chow-fed mice identified *Bacteroides thetaiotaomicron (Bt)* to associate with a subset of cecal lipid metabolites in mice that received PAHSA treatment (Figure 1G). Since a subset of these lipids correlated with insulin sensitivity, we aimed to determine the effects of heat-killed (HKBT) and live (LBT) *Bt* supplementation on gut microbiome structure and function in HFD-fed conventional mice. Principal Components Analysis (PCA) of gut metagenome sequencing data showed that HFD feeding had the greatest effect on gut microbial community composition in both female and male conventional mice compared to effects of treatment with either heat-killed or live *Bt* (Figures 4A-B, Figures S4A-B). Within HFD-fed female mice, microbial composition was slightly more variable in LBT*-*supplemented HFD-female mice compared to HFD-PBS and HFD-HKBT-treated controls (Figure 4A). There was minimal effect of *Bt* supplementation on gut microbiota composition in HFD-fed male mice (Figure 4B).

**Figure 4.**
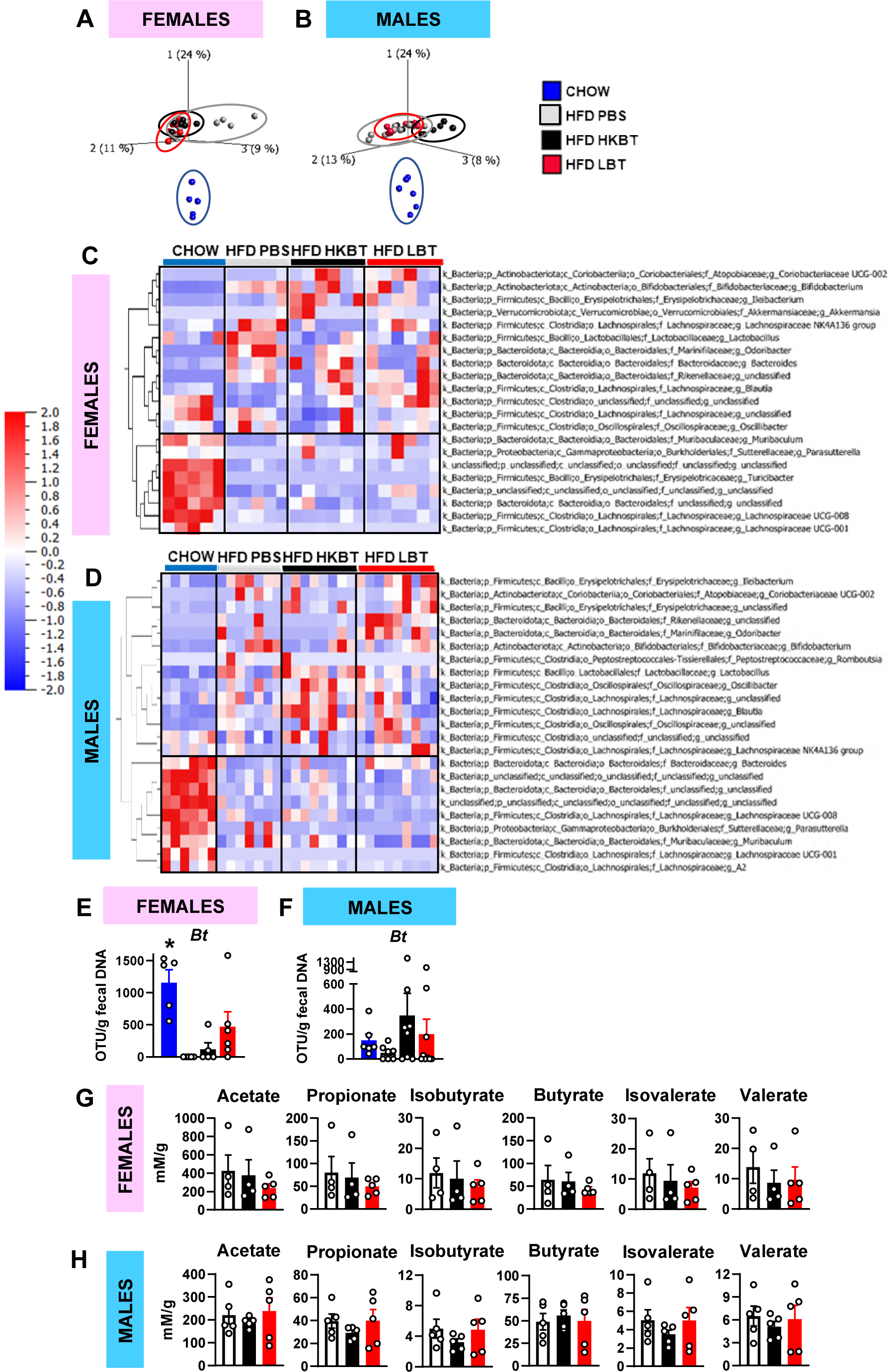
The gut microbiome has sex-specific responses to *Bacteroides thetaiotaomicron* supplementation in dietary obese mice. Conventional C57BL6/J mice were fed chow or HFD supplemented with PBS, heat-killed *Bt* (HKBT), or live *Bt* (LBT) and cecal contents were collected for gut metagenome sequencing. Principal Components Analyses illustrating similarities in the way HFD alters gut microbial composition in **(A)** female and **(B)** male mice with lesser effects of HKBT or LBT. **(C,D)** Heatmap with hierarchical clustering of gut microbial species in **(C)** female and **(D)** male mice. **(E, F)** Abundance levels of *Bt* in **(E)** female and **(F)** male mice, and **(G,H)** short-chain fatty acid levels in the same cecal contents in **(G)** female and **(H)** male mice. For **(A,C,E)** female mice, n=5-6/group. For **(B,D,F)** male mice, n=6-9/group. For **(G-H)** n=4-5/group. *p<0.05 chow vs all groups. All data presented as means ± SEM. Statistical analysis conducted using one-way ANOVA followed by Tukey’s post-hoc test.

Supervised classification and hierarchical clustering of gut microbial taxa demonstrated that female (Figure 4C, Figure S4A) and male (Figure 4D, Figure S4B) mice have distinct sex-specific responses to HFD feeding and *Bt* supplementation. Female chow-fed mice had elevated genera levels of *Turicibacter, Muribaculum* and *Lachnospiraceae (UCG-008* and *UCG-001)* versus all HFD-fed mice (Figure 4C). HFD-PBS female mice had increased genera levels of *Lactobacillus* and *Odoribacter* compared to all other groups (Figure 4C). Although LBT and HKBT had minimal effects on microbial composition, there were some sex specific effects at the genera level. HFD-HKBT female mice had raised genera levels of *Bifidobacterium* and *Coriobacteriaceae* versus chow and HFD-PBS mice, while LBT treatment restored levels of *Blautia* (genera), *Rikenellaceae* (family), and *Clostridia* (class) towards chow levels (Figure 4C). Interestingly, *Blautia* (genera) abundance changed with *Bt* treatment, with levels increasing with HKBT and reaching highest levels with LBT supplementation compared to chow and HFD-PBS controls (Figure 4C).

In contrast, male chow-fed mice had elevated genera levels of *Bacteroides, Parasutterella, Muribaculum,* and *Lachnospiraceae* versus HFD-fed mice (Figure 4D). Within HFD-fed male mice, HKBT treatment, and LBT to a lesser degree, increased genera levels of *Blautia, Lactobacillus,* and *Oscillobacter* compared to PBS controls. HFD-LBT male mice had increased levels of genera that included *Ileibacterium, Coriobacteriaceae (UCG-002),* and *Odoribacter* versus all other groups. These data show that the gut microbiota has sex-specific responses to *Bt* oral bacteriotherapy in HFD-fed mice.

Next, we determined the effect of HFD feeding and *Bt* supplementation on the biomass of detectable *Bt species in vivo*. Metagenome sequencing detected levels of both heat-killed and live *Bt.* HFD-feeding reduced *Bt* levels in female (Figure 4E) and tended to decrease *Bt* levels in male (Figure 4F) mice compared to chow control levels, and both HKBT and LBT supplementation tended to raise *Bt* levels towards those of chow-fed mice. Lastly, we measured gut microbe-derived short-chain fatty acids (SCFAs) but found no effect of HKBT or LBT supplementation on cecal SCFA levels in HFD-fed female (Figure 4G) or in male (Figure 4H) mice.

### *Bacteroides thetaiotaomicron* supplementation improves host metabolism in female dietary obese mice

We next tested the effects of *Bt* supplementation on host metabolism in conventional female and male mice. LBT supplementation decreased weight gain and adiposity in HFD-fed female mice versus HFD-PBS controls (Figures 5A-C). There was no change in lean mass and the decreased adiposity could not be explained by a change in food intake (Figures 5D-E). HFD-fed mice receiving either heat-killed (HKBT) or LBT supplementation tended to have lower ambient glycemia compared to HFD-PBS control mice throughout the course of intervention (Figure 5F). Similarly, glucose tolerance (Figures 5G-H) was improved in HFD-HKBT and HFD-LBT female mice, and this was independent of insulin secretion (Figure 5H). Female LBT-treated mice had increased GLP-1 levels in response to oral lipids versus HFD-PBS controls (Figure 5I). However, *Bt* supplementation had no effect on insulin sensitivity (Figure 5J). The reduced weight gain and adiposity in HFD-LBT mice was not associated with a change in brown adipose tissue *ucp1* expression (Figure 5K). Adipocyte number in perigonadal and subcutaneous white adipose tissues was too variable to detect a statistically significant change and there did not appear to be a difference in adipocyte size (Figure 5L). Fecal energy content was increased in HKBT-treated mice but unchanged in LBT-treated mice compared to controls (Figure 5M). In males fed a HFD, there was no effect of *Bt* administration on any of these metabolic outcomes (Figure S5U-AE). In females fed a chow diet (Fig S5A-J), only the effects of *Bt* on lowering adiposity compared to PBS controls were observed. In males fed a chow diet, *Bt* supplementation had no effect on body weight, body composition, food intake, or insulin sensitivity, but lowered serial glycemia, improved glucose tolerance, and increased glucose-stimulated insulin secretion (5 minutes post oral glucose) (Figures S5K-T).

**Figure 5.**
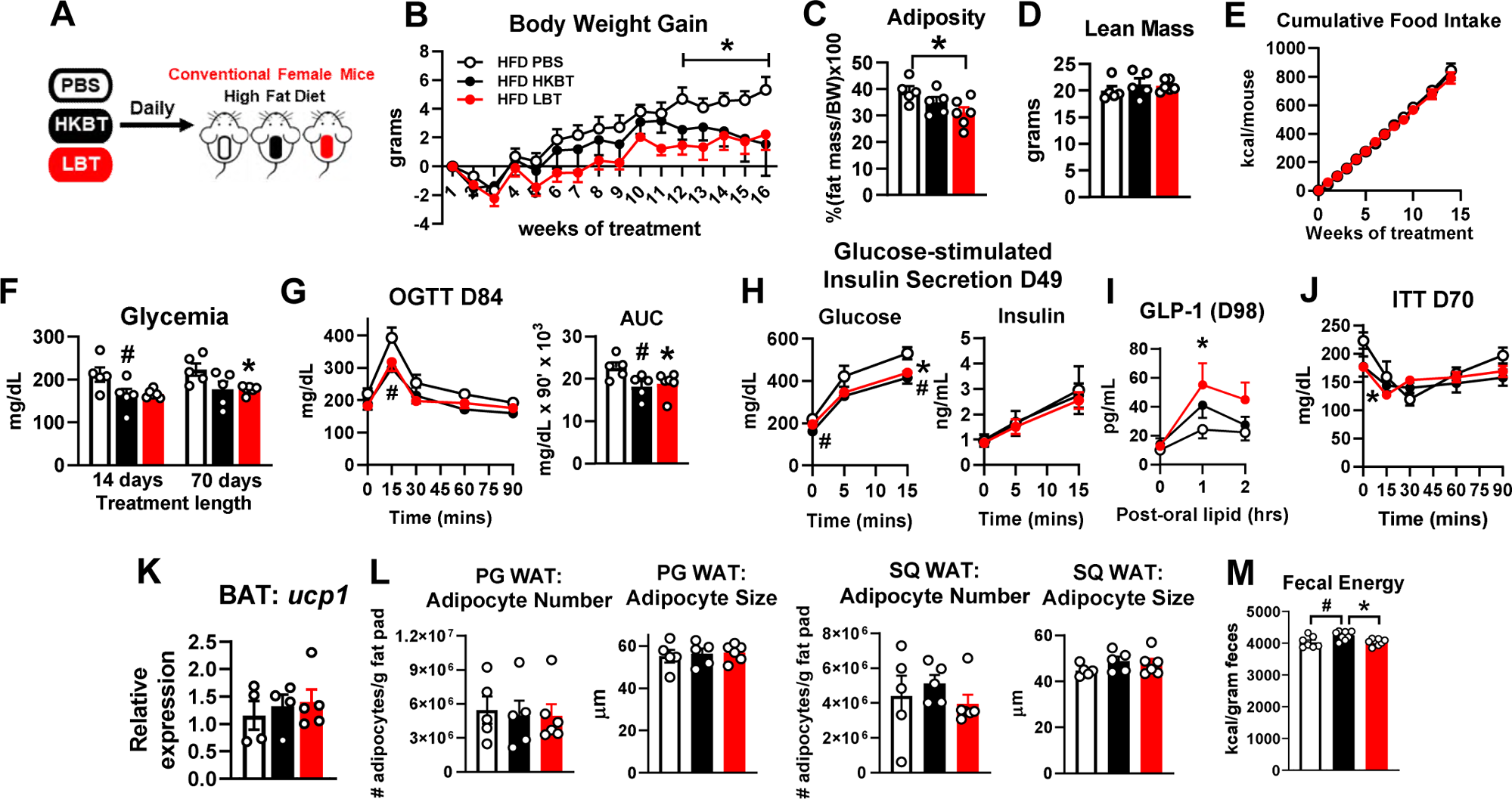
*Bt* supplementation improves host metabolism in conventional diet-induced obese mice. **(A)** Experimental design in which conventional HFD-fed female mice were orally supplemented once daily with PBS, heat-killed *Bt* (HKBT), or live *Bt* (LBT). Effects of *Bt* supplementation on **(B)** weight gain, **(C)** adiposity (110 days of treatment), **(D)** lean mass (110 days of treatment), **(E)** cumulative food intake, **(F)** ambient glycemia following a 5-hour food removal (14 and 70 days of treatment), **(G)** oral glucose tolerance test with area under the curve (AUC) (84 days of treatment), **(H)** circulating glucose and insulin in response to oral glucose (49 days of treatment), **(I)** plasma GLP-1 levels in response to oral lipid (98 days of treatment), **(J)** insulin sensitivity (70 days of treatment), **(K)** *ucp1* expression in brown adipose tissue (BAT), **(L)** adipocyte number and size in perigonadal (PG) and subcutaneous (SC) white adipose tissues (WAT), and **(M)** fecal energy (77 days of treatment). n=5-7 per group. *p<0.05 HFD LBT vs HFD PBS; #p<0.05 HFD HKBT vs HFD PBS. All data presented as means ± SEM. Statistical analysis conducted using one-way ANOVA or repeated measures one-way ANOVA followed by Tukey’s post-hoc test.

Considering the sex specific *Bt* supplementation effects on host metabolism in HFD-fed mice, we asked whether sex hormones may be involved (Figure S5AF). In another cohort of mixed-strain background female mice fed a HFD, LBT supplementation reduced weight gain (Figure S5AG), which is consistent with LBT effects on body weight in pure C57BL6 female mice (Figure 5B). As expected, ovariectomized (OVX) mice gained more weight, tended to have increased adiposity and worsened glucose tolerance (Figures S5AG-AJ) compared to sham-operated controls. HKBT- and LBT-treated HFD-OVX female mice tended to gain less weight and adiposity and were more glucose tolerant compared to HFD-OVX controls (Figures S5AG-AJ), indicating that estrogen only partially mediates the *Bt* effects on host metabolism in dietary obese female mice. Altogether, these data demonstrate that supplementation with live and/or heat-killed *Bt* has beneficial sex-specific effects on host metabolism in conventional dietary obese mice.

### *Bt* supplementation affects the gut mucosa and intestinal innate immune cells in dietary obese mice

Intestinal barrier function plays a critical role in regulating glucose metabolism and is regulated in part by the gut mucosa and gut inflammatory status. HFD feeding decreased colonic mucus thickness in female mice compared to chow-fed controls (Figure 6A). LBT supplementation increased colonic mucus thickness towards those of chow-fed controls (Figure 6A). Similarly, levels of colonic MUC2, the predominant intestinal mucin protein, tended to decrease with HFD feeding, and were ∼70% reduced compared to levels with LBT supplementation in mice on the same diet (Figure 6B). MUC2 is synthesized and secreted by Goblet cells that line the intestinal epithelium. Thus, we measured the number of Goblet cells in ileum from the same mice. Goblet cell frequency was not different between chow and HFD-PBS or HFD-HKBT-treated mice (Figure 6C). However, LBT supplementation increased the number of ileal Goblet cells per villi when compared to HFD-PBS control mice (Figure 6C). By contrast, there was no effect of *Bt* supplementation on the intestinal mucosa in HFD-fed male mice (Figure S6A-B), which may explain the lack of improvement in host metabolism (Figure S5U-AD).

**Figure 6.**
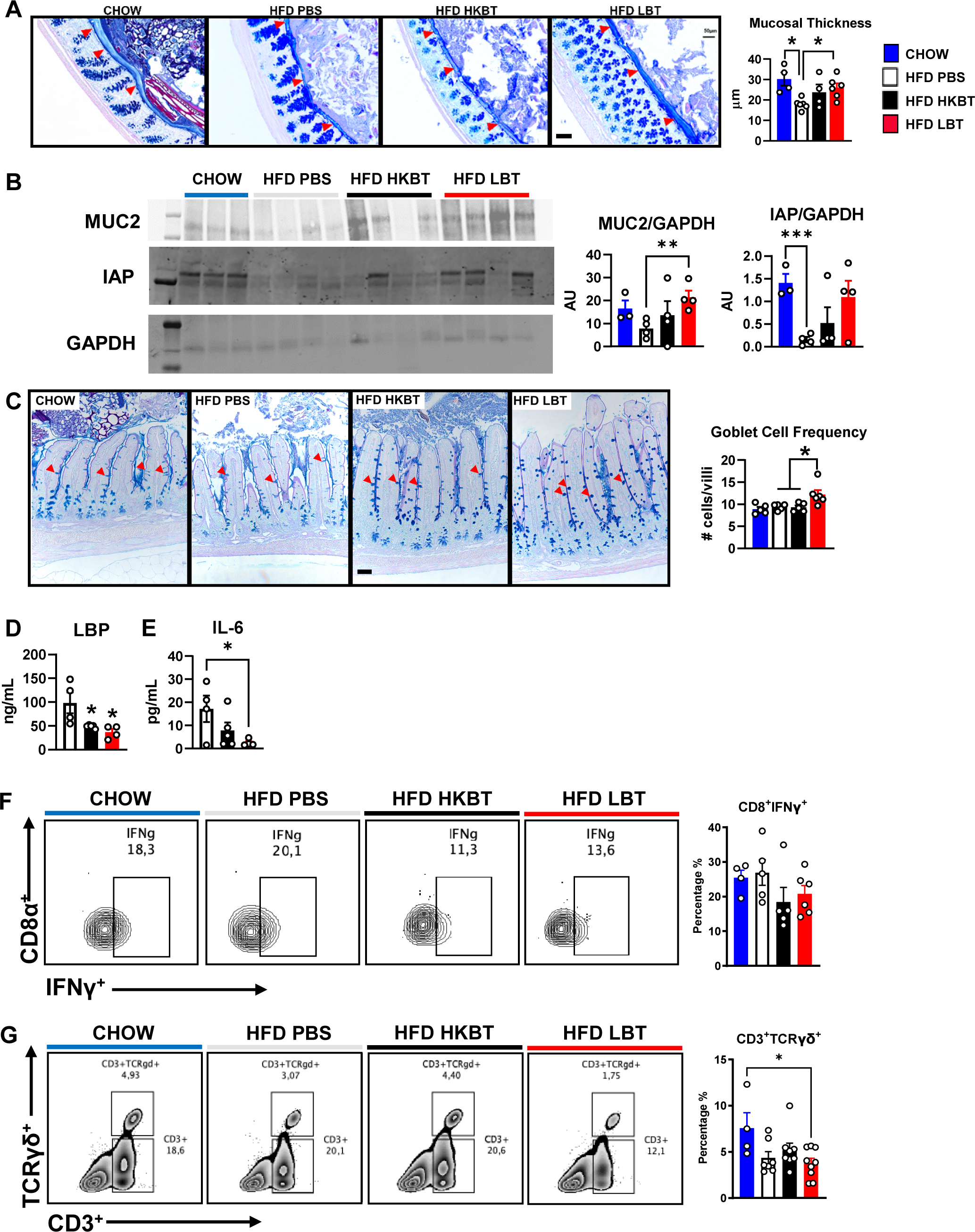
Effects of *Bt* supplementation on the gut mucosa and intestinal innate immune cells in dietary obese mice. Effects of *Bt* supplementation in chow or HFD-fed conventional female mice supplemented with PBS, HKBT or LBT on **(A)** colon mucus thickness, indicated by red arrows (n=4-6/group), **(B)** Western blots and corresponding bar graphs quantifying protein levels of MUC2 (mucin2), IAP (intestinal alkaline phosphatase), and GAPDH (glyceraldehyde 3-phosphate dehydrogenase) by densitometry in mouse ileum (n=3-4/group), **(C)** Goblet cell frequency in mouse ileum, indicated by red arrows, and corresponding bar graph (n=5/group), **(D)** circulating LPS-binding protein ([LBP]; n=4-5/group), **(E)** circulating interleukin-6 (IL-6; n=4-5/group). Intraepithelial lymphocytes (IELs) were isolated from colon, and the percentages of **(F)** CD8^+^IFNγ^+^ cells and **(G)** CD3^+^TCRγδ^+^ cells were measured by flow cytometry (n=4-9/group). All images taken at 20X magnification by light microscopy. Scale bar indicated by the black line = 50µm. *p<0.05 HFD HKBT or HFD LBT vs. HFD PBS, or as indicated. Statistical analysis conducted using one- or two-way ANOVA followed by Tukey’s post-hoc test. All data presented as means ± SEM.

We next measured plasma lipopolysaccharide (LPS) binding protein (LBP), a proxy for LPS levels, and interleukin-6 (IL-6) from the same mice to determine whether the increase in gut mucosa architecture with LBT treatment prevents translocation of pro-inflammatory molecules into systemic circulation. Plasma LBP levels were reduced in both HKBT- and LBT-treated HFD-fed female mice (Figure 6D). Similarly, LBT lowered and HKBT tended to reduce circulating IL-6 levels in dietary obese female mice (Figure 6E). Despite no change in mucosal thickness or Goblet cell frequency, *Bt* supplementation tended to lower levels of circulating LBP and reduced IL-6 in HFD-fed male mice (Figure S6C-D). Altogether, these data indicate that LBT has sex-specific effects to fortify the intestinal mucosa and this associates with a reduction in pro-inflammatory molecules that are known to translocate into systemic circulation in dietary obese mice.

Intestinal innate immune cell types play a critical role in regulating gut homeostasis and contribute to host metabolic processes (5). We find that HFD feeding increases the percentage of colonic intraepithelial CD8+ T-cells compared to chow controls (Figure S6E), and within this population, *Bt* supplementation tended to lower the percentage and MFI of CD8^+^IFNγ^+^ T-cells versus HFD-PBS and chow controls (Figures 6F, S6E). Increasing evidence supports a role for TCRγδ^+^ cells in maintaining gut barrier function, in regulating gut and host metabolism. In our study, HFD feeding tended to lower the percentage of colonic intraepithelial TCRγδ^+^ T-cells compared to chow (Fig 6G), with LBT treatment significantly reducing TCRγδ^+^ levels, although the subset of TCRγδ^+^IFNγ^+^ T-cells was not altered (Fig S6F). Similarly, *Bt* supplementation reduced the expression of T-cell and myeloid cell markers (*tbx21* and *arg1),* and related pro-inflammatory cytokines (*il-17, il-10, tnf-α)* versus HFD-PBS controls (Figure S6H). Overall, these data indicate that *Bt* supplementation modulates colonic intraepithelial immune cell profiles and function.

## DISCUSSION

The gut microbiota offers therapeutic opportunities to improve host metabolism. Here, we demonstrate how anti-diabetic effects of PAHSAs depend on, and can be transmitted by, the gut microbiota. In multi-omic analyses of cecal microbial communities and lipid metabolites in mice with increased insulin sensitivity resulting from PAHSA treatment, we identified *Bacteroides thetaiotamicron* (*Bt*) as a target species associated with beneficial effects on host metabolism. We then treated diet-induced obese mice with *Bt* and found distinct sex-specific effects on the gut microbiome, intestinal barrier function, adiposity, and glucose metabolism. The studies highlight the therapeutic value of targeting the gut microbiota and demonstrate that beneficial effects of the gut microbiota on host metabolism can be sex specific.

Fecal microbiota transplantation (FMT) can transmit host phenotype (11, 29), making FMT an attractive strategy to treat and/or prevent certain human diseases. For example, FMT using healthy donor stool can improve the clinical response of some patients with recurrent *Clostridiodes difficile* infections and irritable bowel disease (IBD) (30–32), although effects are variable with rare cases of complications (33, 34). Despite the known risks, interest remains in applying FMT for the treatment and/or prevention of obesity and associated metabolic diseases (35–40). We find that the beneficial PAHSA effects to improve insulin sensitivity in mice are transmissible by FMT. Donor feces from male mice resulted in increased insulin sensitivity in male, but not female, recipient mice, highlighting the importance of sex in mediating gut microbiome effects on host metabolism (41, 42). Overall, our studies demonstrate that the gut microbiota plays a key role in mediating PAHSA effects to improve glucose metabolism and support the broader notion that the gut microbiome is a therapeutic target that may be leveraged to improve host metabolism in humans with obesity.

Phenotype transmission by FMT led us to ask whether a specific microbiota contributed to the beneficial PAHSA effects on glucose metabolism. We are the first to report the effects of PAHSA treatment on gut microbiota composition in mice. Functional analysis showed that *Bt* positively associated with a signature of lipid species in mice with increased insulin sensitivity in response to PAHSA treatment. *Bt* levels were not increased with PAHSA treatment, but it is possible that *Bt* may have functional effects on host metabolism because the function of gut microbiota, including *Bt,* depends on substrate availability and the intestinal microenvironment (43).

This led us to test whether *Bt* supplementation has beneficial effects to improve host metabolism in diet-induced obese mice. We found that 16 weeks of daily oral *Bt* supplementation in female mice fed a HFD for 28 weeks starting at 24 weeks of age (52-weeks-old at terminal tissue collection) restored *Bt* levels towards those of chow control mice, reduced weight gain and adiposity, and improved glucose tolerance. This could be relevant to human obesity since *Bt* levels are low in humans with obesity and restored with bariatric surgery. In our studies, *Bt* supplementation had no effect on host metabolism in HFD-fed male mice. This sex-specific response to *Bt* supplementation may be due, in part, to inherent differences in the gut microbiotas of male and female C57BL6 mice (44). One study in 15-week-old HFD-fed male mice showed that 7 weeks of *Bt* supplementation reduced weight gain, adiposity, and adipocyte size (45). In another study, 3 weeks of *Bt* supplementation to 6-week-old C57BL6 male mice fed a HFD for 3 weeks improved insulin sensitivity (46). The different outcomes between our study and others may be due to differences in mouse age, type of HFD (Td93073, Research Diets D12492i, QuickFat CLEAJapan), mouse vendor sources, treatment protocols, and housing conditions across experimental animal facilities, all of which can impact gut microbiome-host outcomes (47, 48). Also, in our studies, we administered *Bt* daily, whereas *Bt* was administered thrice weekly in other studies (45, 46). The fact that *Bt* was identified from chow-fed male mice treated with PAHSAs, but *Bt* supplementation did not improve host metabolism in HFD-fed male mice, whereas FMT did, suggests that other gut microbiota may also play a role in improving metabolism in HFD-fed male mice. The fact that *Bt* supplementation reduced weight gain in female mice fed a HFD but HFD-fed germ-free female mice did not respond to FMT may reflect lower abundance levels of *Bt* with FMT compared to *Bt* supplementation. This may also reflect differences in how conventional and germ-free female and male mice respond to FMT and *Bt* supplementation. Overall, our findings suggest that *Bt* supplementation in female mice with dietary-induced obesity may improve several metabolic parameters.

Sex hormones regulate the function of metabolic tissues (49) and contribute to differences in gut microbiome composition (41). However, the role of sex hormones in mediating probiotic supplementation effects on host metabolism is less known. To address this knowledge gap and based on our result that *Bt* supplementation improved several metabolic parameters only in HFD-fed female mice, we performed ovariectomy studies in diet-induced obese mice. We found that ovariectomy partially reversed some of the *Bt* effects on host metabolism. Many commercial probiotics are marketed to target specific demographics (infant, women, all adults) for specific health outcomes (for example, immune health, bloating, diarrhea). Most open formulations comprise common subsets of *Lactobacilli* and *Bifidobacterium* strains differing only in bacterial concentration. There is a growing market of next-generation probiotics to target metabolism and weight management, but their formulations are largely proprietary and effects in both sexes are less clear (50). Our *Bt* supplementation and ovariectomy studies are among the few to date determining the role of estrogen and sex as a biological variable in contributing to probiotic effects on host metabolism. Our work highlights the need for more studies to determine sex-specific responses to probiotic supplementation.

We observed that both heat-killed and live *Bt* had comparable effects to reduce weight gain, improve glucose tolerance, and lower circulating LPS and IL-6 in HFD-fed female mice. Both live and pasteurized (heat-killed) probiotics have therapeutic efficacy to improve multiple metabolic parameters in mice and in humans (51, 52). For example, live and pasteurized *Akkermansia muciniphila* supplementation reduced weight gain and improved insulin sensitivity in mice and in proof-of-concept human studies (50–52). In one study, pasteurized *Akkermansia muciniphila* retained its ability to bind to and activate TLR2, and this improved host metabolism in mice (52). Increasing the shelf stability of probiotics by pasteurization will make them more practical to use. Thus, it is advantageous that *Bt* is effective even after pasteurization. Future studies are needed to determine the mechanism by which pasteurized *Bt* acts to improve host metabolism in mice.

Fortifying the intestinal barrier has therapeutic potential to prevent and/or treat metabolic disease (6, 13, 53). HFD-induced obesity increases gut permeability and systemic inflammation, which worsens host metabolism in mice (54). These effects are also observed in humans (16). We report that despite HFD feeding in female mice, live *Bt* supplementation preserves colon mucus thickness, and this is associated with increased mucin protein levels and Goblet cell frequency to levels of chow-fed control mice. This result is consistent with studies in which *Akkermansia muciniphila* supplementation increased Goblet cell density in the ileum of diet-induced obese mice (52), and germ-free rats supplemented with *Bt* had increased Goblet cell differentiation and elevated expression of mucus-related genes (55). We also report that *Bt* supplementation increases protein levels of intestinal alkaline phosphatase (IAP), the primary LPS-detoxifying enzyme that resides within the intestinal brush border membrane (56, 57). We show that the reduced weight gain and adiposity in HFD-fed mice supplemented with live *Bt* is associated with elevated colon IAP levels, which may reflect a proportional increase in mucus thickness. This result is consistent with studies in HFD-fed mice in which IAP inhibition raised circulating levels of LBP and TNF-α (44), and in human studies in which people with Type 2 Diabetes Mellitus have lower stool IAP levels compared to people without diabetes (58). Overall, we show that *Bt* supplementation fortifies the gut mucus barrier, which may contribute to preventing the translocation of pro-inflammatory gut microbe derived LPS from entering host circulation.

We also find that colonic IEL T-cells play a minimal role in mediating *Bt* effects on host metabolism in HFD-fed female mice. This may be due to the lack of change in levels of fecal short-chain fatty acids (SCFAs), which are critical regulators of T-cell function (59, 60). Mucosal T-cells play an important role in regulating intestinal epithelial cell turnover and repair to maintain gut barrier function (61, 62). Mucosal T-cells expressing TCRγδ reside predominantly within the mucosal epithelium to sustain gut barrier integrity. Here, HFD feeding tended to lower colonic IEL TCRγδ^+^ levels, but *Bt* supplementation was not sufficient to restore these levels towards chow controls. Previous studies demonstrate that different T-cell sub-types (Th1, Th17, Tregs) are critical to promote colonic homeostasis and health in mice (5, 59), and therapeutic targeting of specific T-cell sub-types is sufficient to delay HFD-induced senescence in mice (63). These studies measured T-cell subtypes in the colonic lamina propria in HFD-fed male mice, while our studies measured intraepithelial T-cell subtypes in female mice, which may explain the reported differences in mucosal T-cell responses to HFD feeding in mice. Overall, our data indicate that *Bt* supplementation has some beneficial effects on the intestinal mucosa but is independent of mucosal T-cells in HFD-fed female mice.

We present the first analyses of PAHSA treatment effects on plasma and cecal lipid metabolite profiles in mice. PAHSAs differentially regulated levels of multiple lipid species, including cholesterol esters (CEs), phosphatidylethanolamines (PEs), and triacylglycerides (TAGs). Furthermore, the levels of PE species correlated with *Bt*. This could have therapeutic implications, since levels of PE, the predominant phospholipid in the body and a lipid precursor for phosphatidylcholine (PC), have been shown to be associated with insulin sensitivity, BMI, and waist-to-hip ratio (WHR) in humans with obesity (64, 65). Our data suggest that PAHSA treatment may regulate levels of some PE species, and this may contribute to the enhanced insulin sensitivity observed in mice, although the studies here did not test this directly. Overall, our results identify new lipid species and signatures in mice with increased insulin sensitivity resulting from PAHSA treatment. Future studies are needed to determine the therapeutic efficacy of these lipid metabolites in the context of diet-induced obesity.

Overall, the studies presented here highlight the therapeutic potential of targeting the gut microbiome to beneficially modulate host metabolism and prevent obesity. These studies were performed in both sexes of mice and highlight the critical role sex dimorphism plays in regulating energy balance and host metabolism. Identifying novel gut microbiota and optimizing next-generation microbe-based interventions have major potential to complement existing therapies and/or strategies to support long-term weight and glucose management in people with obesity. Increasing our knowledge of how gut microbiota functions in response to lipids, including PAHSAs, may unlock new therapeutic targets and strategies that leverage the gut microbiome to improve host health.

### Limitations of Study

In the FMT studies reported here, donor feces from PAHSA-treated male mice transmitted some of the beneficial effects of PAHSAs on glucose metabolism to recipient GF male, but not female, mice. Whether donor feces from female PAHSA-treated mice can transmit metabolic effects to recipient GF female mice requires further investigation. However, we can still conclude that some of the beneficial effects of PAHSAs on glucose metabolism can be transmitted by FMT. It is also important to note that applying b-test for multi-omics functional analysis of the cecal metagenome and metabolome identified *Bt* and several lipid species that associated with mice that were insulin sensitive in response to PAHSA treatment. Future studies are needed to determine the effects of supplementation with these lipid metabolites on improving host metabolism in the setting of diet-induced obesity.

## AUTHOR CONTRIBUTIONS

J.L. conceived of, designed, performed, and interpreted experiments and data. B.B.K. supervised the experimental plan and interpreted experiments and data. K.W. assisted with animal studies and performed immunology experiments. A.T.N. and D.S. performed chemical synthesis of PAHSA lipids. V.L. and L.B. assisted with and interpreted data related to gnotobiotic mouse studies. M.M.H. and B.P.C designed and measured GLP-1 assays. A.C. designed and interpreted immunology experiments. A.R. developed and performed multi-omics analyses. C.C. performed lipidomics measurements. J.L and B.B.K. wrote the manuscript.

## ACKNOWLEDGMENTS

The Harvard Digestive Disease Center and NIH grant P30 DK034854 supported the Histology and Imaging Core. P30 DK034854 and a capital grant from the Massachusetts Life Sciences Center supported gnotobiotic mouse studies and microbial preparation of *Bacteroides thetaiotaomicron* by the Massachusetts Host Microbiome Center. We thank Dr. Yana Stackpole for providing bioinformatics support, and Mary Louise Delaney, MA, for microbiology support.

## FUNDING SOURCES

1K01DK114162-01A1 (JL), BADERC 5P30DK057521-19 (JL), HDDC P30 DK034854 (JL), JPB (BBK) and R01 DK106210 (BBK).

---

**Figure S1, related to Figure 1. PAHSA treatment in mice increases insulin sensitivity and alters the gut microbiota and plasma and cecal metabolomes.**

(A) Gene expression of *gcg* and *clca1* in ileum from chow-fed male mice which were treated with oral PAHSAs for 17 days. Samples were collected after mice were fasted overnight and refed for 5-hours. (B) Gene expression of pro-inflammatory T-cell markers and related cytokines and *occludin* in ileum from HFD-fed male mice treated with 9-PAHSA by subcutaneous minipumps for 6 months. Samples were collected in the ad lib fed state. n=6-8/group. *p<0.05 vehicle vs PAHSA. Statistical analysis conducted by Student’s t-test. All data presented as means ± SEM.

**Figure S2, related to Figure 2. The beneficial PAHSA effects on glucose homeostasis are transmissible by fecal microbiota transplantation.**

**(A)** Experimental design for fecal microbiota transplantation (FMT) studies in which feces from donor chow-fed male mice treated with vehicle (V) or PAHSAs (P) were orally administered to recipient HFD-fed germ-free (GF) female mice. GF-HFD mice that did not receive FMT (HFD no FMT) served as controls. FMT effects in recipient GF-HFD female mice on **(B)** body weight, **(C)** weight gain, **(D)** intraperitoneal glucose tolerance 21 days post-FMT, **(E)** oral glucose tolerance 28 days post-FMT, and **(F)** insulin sensitivity 35 days post-FMT. n=5 GF-HFD mice not receiving FMT (No FMT), n=8. GF-HFD mice receiving FMT from VEH-treated mice, n=8 GF-HFD mice receiving FMT from PAHSA-treated mice.

**Figure S3, related to Figure 3. The gut microbiota is necessary for PAHSA to improve glucose homeostasis in HFD-fed mice.**

**(A)** Bacterial culture plates using terminal fecal pellets collected from gnotobiotic isolators housing experimental GF-HFD mice treated with vehicle or PAHSAs confirms gnotobiotic status during treatment intervention.

**Figure S4, related to Figure 4. The gut microbiome has sex-specific responses to *Bt* supplementation in dietary obese mice.**

Heatmap with hierarchical clustering of all gut microbial species from gut metagenome sequencing of cecal contents from **(A)** female and **(B)** male conventional mice fed chow or HFD supplemented with PBS, heat-killed *Bt* (HKBT), or live *Bt* (LBT).

**Figure S5, related to Figure 5. *Bt* supplementation improves host metabolism in conventional diet-induced obese mice.**

**(A)** Experimental design in which conventional female chow-fed mice were orally supplemented with PBS, heat-killed *Bt* (HKBT), or live *Bt* (LBT) three times per week. Effects of *Bt* supplementation on **(B)** weight gain, **(C)** adiposity (60 days of treatment), **(D)** lean mass (60 days of treatment), **(E)** cumulative food intake, **(F)** ambient glycemia following a 5-hour food removal, **(G)** intraperitoneal glucose tolerance test (IPGTT; 42 days of treatment), **(H)** oral glucose tolerance test (37 days of treatment), **(I)** glucose and insulin levels during an oral glucose-stimulated insulin secretion assay in mice (56 days of treatment), **(J)** insulin sensitivity (49 days of treatment). For **(A-J):** n=6-7/group. *p<0.05 LBT vs PBS, #p<0.05 HKBT vs PBS, +p<0.05 LBT vs HKBT, determined by one-way ANOVA followed by Tukey’s post-hoc test. All data presented as means ± SEM. **(K)** Experimental design where conventional female chow-fed mice were orally supplemented with PBS, heat-killed *Bt* (HKBT), or live *Bt* (LBT) three times per week. Effects of *Bt* supplementation on **(L)** weight gain, **(M)** adiposity (60 days of treatment), **(N)** lean mass (60 days of treatment), **(O)** cumulative food intake, **(P)** ambient glycemia following a 5-hour food removal, **(Q)** intraperitoneal glucose tolerance test (IPGTT; 42 days of treatment), **(R)** oral glucose tolerance test (OGTT; 37 days of treatment), **(S)** glucose and insulin levels during an oral glucose-stimulated insulin secretion assay in mice (56 days of treatment), and **(T)** insulin sensitivity (49 days of treatment). For **(K-T):** n=6-7/group. *p<0.05 LBT vs PBS; #p<0.05 HKBT vs PBS; +p<0.05 LBT vs HKBT, determined by two-way ANOVA followed by Tukey’s post-hoc test. All data presented as means ± SEM. **(U)** Experimental design in which conventional HFD-fed male mice were orally supplemented with PBS, heat-killed *Bt* (HKBT), or live *Bt* (LBT) once daily. Effects of *Bt* supplementation on: **(V)** weight gain, **(W)** adiposity (110 days of treatment), **(X)** lean mass (110 days of treatment), **(Y)** cumulative food intake, **(Z)** ambient glycemia following a 5-hour food removal, **(AA)** oral glucose tolerance test (OGTT; 91 days of treatment), **(AB)** glucose and insulin levels during an oral glucose tolerance test (49 days of treatment), **(AC)** plasma GLP-1 levels in response to oral lipid (98 days of treatment), **(AD)** insulin sensitivity (77 days of treatment), **(AE)** Fecal energy (110 days of treatment). For **(U-AE):** n=5-9/group. All data presented as means ± SEM. **(AF)** Experimental design in which conventional HFD-fed female mice received sham or ovariectomy surgery and were orally supplemented with PBS, HKBT or LBT. Effects of *Bt* supplementation on **(AG)** weight gain, **(AH)** adiposity (47 days of treatment), **(AI)** area under the curve (AUC) representing glucose tolerance (25 days of treatment), and **(AJ)** area under the curve (AUC) for glucose and insulin levels during an oral glucose-stimulated insulin secretion assay in mice (50 days of treatment). For **(AF-AJ):** *p<0.05 as indicated. N=5-6/group. All data presented as means ± SEM.

**Figure S6, related to Figure 6. Effects of *Bt* supplementation on the gut mucosa and intestinal innate immune cells in dietary obese mice.**

Effects of *Bt* supplementation in chow or HFD-fed conventional male mice supplemented with PBS, HKBT or LBT on **(A)** colon mucus thickness (n=2-4/group), **(B)** Goblet cell frequency in mouse ileum (n=5-8/group), **(C)** circulating LPS-binding protein (LBP; n=4-5/group), and **(D)** circulating interleukin-6 (IL-6; n=5-7/group). For **(A-B):** All histological images taken at 10X magnification by light microscopy. Scale bar indicated by the black line = 100µm. Intraepithelial lymphocytes (IELs) were isolated from colon from HFD-fed conventional female mice, and the percentages of **(E)** CD8^+^ and CD8^+^IFNɤ^+^, and **(F)** TCRɤδ^+^IFNɤ^+^ cells were measured by flow cytometry (n=3-9/group). **(G)** Gating strategy used to identify CD3^+^TCRɤδ^+^ and CD8^+^IFNɤ^+^ colonic IELs. Expression of genes for innate and adaptive immune cells and related cytokines in HFD-fed conventional female mice supplemented with PBS, HKBT or LBT (n=2-5/group). *p<0.05 HFD LBT or HFD HKBT vs HFD PBS; #p<0.05 HFD PBS or HFD HKBT vs chow. Statistical analysis conducted using one-way ANOVA followed by Tukey’s post-hoc test. All data presented as means ± SEM.

## Notes

### Competing Interest Statement

The authors have declared no competing interest.

## REFERENCES

1. (WHO) WHO. Obesity and overweight 2021. Available from: https://www.who.int/news-room/fact-sheets/detail/obesity-and-overweight.

2. Jaacks LM, Vandevijvere S, Pan A, McGowan CJ, Wallace C, Imamura F, Mozaffarian D, Swinburn B, Ezzati M. The obesity transition: stages of the global epidemic. Lancet Diabetes Endocrinol. 2019;7(3):231–40. Epub 2019/02/02. doi: 10.1016/s2213-8587(19)30026-9. PubMed PMID: 30704950; PMCID: PMC7360432.

3. Afshin A, Forouzanfar MH, Reitsma MB, Sur P, Estep K, Lee A, Marczak L, Mokdad AH, Moradi-Lakeh M, Naghavi M, Salama JS, Vos T, Abate KH, Abbafati C, Ahmed MB, Al-Aly Z, Alkerwi A, Al-Raddadi R, Amare AT, Amberbir A, Amegah AK, Amini E, Amrock SM, Anjana RM, Ärnlöv J, Asayesh H, Banerjee A, Barac A, Baye E, Bennett DA, Beyene AS, Biadgilign S, Biryukov S, Bjertness E, Boneya DJ, Campos-Nonato I, Carrero JJ, Cecilio P, Cercy K, Ciobanu LG, Cornaby L, Damtew SA, Dandona L, Dandona R, Dharmaratne SD, Duncan BB, Eshrati B, Esteghamati A, Feigin VL, Fernandes JC, Fürst T, Gebrehiwot TT, Gold A, Gona PN, Goto A, Habtewold TD, Hadush KT, Hafezi-Nejad N, Hay SI, Horino M, Islami F, Kamal R, Kasaeian A, Katikireddi SV, Kengne AP, Kesavachandran CN, Khader YS, Khang YH, Khubchandani J, Kim D, Kim YJ, Kinfu Y, Kosen S, Ku T, Defo BK, Kumar GA, Larson HJ, Leinsalu M, Liang X, Lim SS, Liu P, Lopez AD, Lozano R, Majeed A, Malekzadeh R, Malta DC, Mazidi M, McAlinden C, McGarvey ST, Mengistu DT, Mensah GA, Mensink GBM, Mezgebe HB, Mirrakhimov EM, Mueller UO, Noubiap JJ, Obermeyer CM, Ogbo FA, Owolabi MO, Patton GC, Pourmalek F, Qorbani M, Rafay A, Rai RK, Ranabhat CL, Reinig N, Safiri S, Salomon JA, Sanabria JR, Santos IS, Sartorius B, Sawhney M, Schmidhuber J, Schutte AE, Schmidt MI, Sepanlou SG, Shamsizadeh M, Sheikhbahaei S, Shin MJ, Shiri R, Shiue I, Roba HS, Silva DAS, Silverberg JI, Singh JA, Stranges S, Swaminathan S, Tabarés-Seisdedos R, Tadese F, Tedla BA, Tegegne BS, Terkawi AS, Thakur JS, Tonelli M, Topor-Madry R, Tyrovolas S, Ukwaja KN, Uthman OA, Vaezghasemi M, Vasankari T, Vlassov VV, Vollset SE, Weiderpass E, Werdecker A, Wesana J, Westerman R, Yano Y, Yonemoto N, Yonga G, Zaidi Z, Zenebe ZM, Zipkin B, Murray CJL. Health Effects of Overweight and Obesity in 195 Countries over 25 Years. N Engl J Med. 2017;377(1):13–27. Epub 2017/06/13. doi: 10.1056/NEJMoa1614362. PubMed PMID: 28604169; PMCID: PMC5477817.

4. Winer DA, Luck H, Tsai S, Winer S. The Intestinal Immune System in Obesity and Insulin Resistance. Cell Metab. 2016;23(3):413–26. Epub 2016/02/09. doi: 10.1016/j.cmet.2016.01.003. PubMed PMID: 26853748.

5. Luck H, Tsai S, Chung J, Clemente-Casares X, Ghazarian M, Revelo XS, Lei H, Luk CT, Shi SY, Surendra A, Copeland JK, Ahn J, Prescott D, Rasmussen BA, Chng MH, Engleman EG, Girardin SE, Lam TK, Croitoru K, Dunn S, Philpott DJ, Guttman DS, Woo M, Winer S, Winer DA. Regulation of obesity-related insulin resistance with gut anti-inflammatory agents. Cell Metab. 2015;21(4):527–42. Epub 2015/04/12. doi: 10.1016/j.cmet.2015.03.001. PubMed PMID: 25863246.

6. Schroeder BO, Birchenough GMH, Ståhlman M, Arike L, Johansson MEV, Hansson GC, Bäckhed F. Bifidobacteria or Fiber Protects against Diet-Induced Microbiota-Mediated Colonic Mucus Deterioration. Cell Host Microbe. 2018;23(1):27–40.e7. Epub 2017/12/26. doi: 10.1016/j.chom.2017.11.004. PubMed PMID: 29276171; PMCID: PMC5764785.

7. Nieuwdorp M, Gilijamse PW, Pai N, Kaplan LM. Role of the microbiome in energy regulation and metabolism. Gastroenterology. 2014;146(6):1525–33. Epub 2014/02/25. doi: 10.1053/j.gastro.2014.02.008. PubMed PMID: 24560870.

8. Cox AJ, West NP, Cripps AW. Obesity, inflammation, and the gut microbiota. Lancet Diabetes Endocrinol. 2015;3(3):207–15. Epub 2014/07/30. doi: 10.1016/s2213-8587(14)70134-2. PubMed PMID: 25066177.

9. Ley RE, Turnbaugh PJ, Klein S, Gordon JI. Microbial ecology: human gut microbes associated with obesity. Nature. 2006;444(7122):1022–3. Epub 2006/12/22. doi: 10.1038/4441022a. PubMed PMID: 17183309.

10. Turnbaugh PJ, Ley RE, Mahowald MA, Magrini V, Mardis ER, Gordon JI. An obesity-associated gut microbiome with increased capacity for energy harvest. Nature. 2006;444(7122):1027–31. Epub 2006/12/22. doi: 10.1038/nature05414. PubMed PMID: 17183312.

11. Ridaura VK, Faith JJ, Rey FE, Cheng J, Duncan AE, Kau AL, Griffin NW, Lombard V, Henrissat B, Bain JR, Muehlbauer MJ, Ilkayeva O, Semenkovich CF, Funai K, Hayashi DK, Lyle BJ, Martini MC, Ursell LK, Clemente JC, Van Treuren W, Walters WA, Knight R, Newgard CB, Heath AC, Gordon JI. Gut microbiota from twins discordant for obesity modulate metabolism in mice. Science. 2013;341(6150):1241214. Epub 2013/09/07. doi: 10.1126/science.1241214. PubMed PMID: 24009397; PMCID: PMC3829625.

12. Turnbaugh PJ, Bäckhed F, Fulton L, Gordon JI. Diet-induced obesity is linked to marked but reversible alterations in the mouse distal gut microbiome. Cell Host Microbe. 2008;3(4):213–23. Epub 2008/04/15. doi: 10.1016/j.chom.2008.02.015. PubMed PMID: 18407065; PMCID: PMC3687783.

13. Cani PD, Possemiers S, Van de Wiele T, Guiot Y, Everard A, Rottier O, Geurts L, Naslain D, Neyrinck A, Lambert DM, Muccioli GG, Delzenne NM. Changes in gut microbiota control inflammation in obese mice through a mechanism involving GLP-2-driven improvement of gut permeability. Gut. 2009;58(8):1091–103. Epub 2009/02/26. doi: 10.1136/gut.2008.165886. PubMed PMID: 19240062; PMCID: PMC2702831.

14. de La Serre CB, Ellis CL, Lee J, Hartman AL, Rutledge JC, Raybould HE. Propensity to high-fat diet-induced obesity in rats is associated with changes in the gut microbiota and gut inflammation. Am J Physiol Gastrointest Liver Physiol. 2010;299(2):G440–8. Epub 2010/05/29. doi: 10.1152/ajpgi.00098.2010. PubMed PMID: 20508158; PMCID: PMC2928532.

15. Desai MS, Seekatz AM, Koropatkin NM, Kamada N, Hickey CA, Wolter M, Pudlo NA, Kitamoto S, Terrapon N, Muller A, Young VB, Henrissat B, Wilmes P, Stappenbeck TS, Núñez G, Martens EC. A Dietary Fiber-Deprived Gut Microbiota Degrades the Colonic Mucus Barrier and Enhances Pathogen Susceptibility. Cell. 2016;167(5):1339–53.e21. Epub 2016/11/20. doi: 10.1016/j.cell.2016.10.043. PubMed PMID: 27863247; PMCID: PMC5131798.

16. Seethaler B, Basrai M, Neyrinck AM, Nazare JA, Walter J, Delzenne NM, Bischoff SC. Biomarkers for assessment of intestinal permeability in clinical practice. Am J Physiol Gastrointest Liver Physiol. 2021;321(1):G11–g7. Epub 2021/05/20. doi: 10.1152/ajpgi.00113.2021. PubMed PMID: 34009040.

17. Aryal P, Syed I, Lee J, Patel R, Nelson AT, Siegel D, Saghatelian A, Kahn BB. Distinct biological activities of isomers from several families of branched fatty acid esters of hydroxy fatty acids (FAHFAs). J Lipid Res. 2021;62:100108. Epub 2021/08/22. doi: 10.1016/j.jlr.2021.100108. PubMed PMID: 34418413; PMCID: PMC8479484.

18. Lee J, Moraes-Vieira PM, Castoldi A, Aryal P, Yee EU, Vickers C, Parnas O, Donaldson CJ, Saghatelian A, Kahn BB. Branched Fatty Acid Esters of Hydroxy Fatty Acids (FAHFAs) Protect against Colitis by Regulating Gut Innate and Adaptive Immune Responses. J Biol Chem. 2016;291(42):22207–17. Epub 2016/08/31. doi: 10.1074/jbc.M115.703835. PubMed PMID: 27573241; PMCID: PMC5064000.

19. Syed I, Lee J, Moraes-Vieira PM, Donaldson CJ, Sontheimer A, Aryal P, Wellenstein K, Kolar MJ, Nelson AT, Siegel D, Mokrosinski J, Farooqi IS, Zhao JJ, Yore MM, Peroni OD, Saghatelian A, Kahn BB. Palmitic Acid Hydroxystearic Acids Activate GPR40, Which Is Involved in Their Beneficial Effects on Glucose Homeostasis. Cell Metab. 2018;27(2):419–27.e4. Epub 2018/02/08. doi: 10.1016/j.cmet.2018.01.001. PubMed PMID: 29414687; PMCID: PMC5807007.

20. Yore MM, Syed I, Moraes-Vieira PM, Zhang T, Herman MA, Homan EA, Patel RT, Lee J, Chen S, Peroni OD, Dhaneshwar AS, Hammarstedt A, Smith U, McGraw TE, Saghatelian A, Kahn BB. Discovery of a class of endogenous mammalian lipids with anti-diabetic and anti-inflammatory effects. Cell. 2014;159(2):318–32. Epub 2014/10/11. doi: 10.1016/j.cell.2014.09.035. PubMed PMID: 25303528; PMCID: PMC4260972.

21. Zhou P, Santoro A, Peroni OD, Nelson AT, Saghatelian A, Siegel D, Kahn BB. PAHSAs enhance hepatic and systemic insulin sensitivity through direct and indirect mechanisms. J Clin Invest. 2019;129(10):4138–50. Epub 2019/08/27. doi: 10.1172/jci127092. PubMed PMID: 31449056; PMCID: PMC6763232.

22. Carvalho E, Kotani K, Peroni OD, Kahn BB. Adipose-specific overexpression of GLUT4 reverses insulin resistance and diabetes in mice lacking GLUT4 selectively in muscle. Am J Physiol Endocrinol Metab. 2005;289(4):E551–61. Epub 2005/06/02. doi: 10.1152/ajpendo.00116.2005. PubMed PMID: 15928024.

23. Shepherd PR, Gnudi L, Tozzo E, Yang H, Leach F, Kahn BB. Adipose cell hyperplasia and enhanced glucose disposal in transgenic mice overexpressing GLUT4 selectively in adipose tissue. J Biol Chem. 1993;268(30):22243–6. Epub 1993/10/25. PubMed PMID: 8226728.

24. Souza VR, Mendes E, Casaro M, Antiorio A, Oliveira FA, Ferreira CM. Description of Ovariectomy Protocol in Mice. Methods Mol Biol. 2019;1916:303–9. Epub 2018/12/12. doi: 10.1007/978-1-4939-8994-2_29. PubMed PMID: 30535707.

25. James OJ, Vandereyken M, Swamy M. Isolation, Characterization, and Culture of Intestinal Intraepithelial Lymphocytes. Methods Mol Biol. 2020;2121:141–52. Epub 2020/03/10. doi: 10.1007/978-1-0716-0338-3_13. PubMed PMID: 32147793.

26. Lab TH. KneadData. Available from: https://github.com/biobakery/kneaddata.

27. Franzosa EA, McIver LJ, Rahnavard G, Thompson LR, Schirmer M, Weingart G, Lipson KS, Knight R, Caporaso JG, Segata N, Huttenhower C. Species-level functional profiling of metagenomes and metatranscriptomes. Nat Methods. 2018;15(11):962–8. Epub 2018/11/01. doi: 10.1038/s41592-018-0176-y. PubMed PMID: 30377376; PMCID: PMC6235447.

28. Bahar Sayoldin MB, Keith A. Crandall, Ali Rahnavard. btest: link, rank, and visualize associations among omics features across multi-omics datasets 2023. Available from: https://github.com/omicsEye/btest.

29. Turnbaugh PJ, Ridaura VK, Faith JJ, Rey FE, Knight R, Gordon JI. The effect of diet on the human gut microbiome: a metagenomic analysis in humanized gnotobiotic mice. Sci Transl Med. 2009;1(6):6ra14. Epub 2010/04/07. doi: 10.1126/scitranslmed.3000322. PubMed PMID: 20368178; PMCID: PMC2894525.

30. El-Salhy M, Hatlebakk JG, Gilja OH, Bråthen Kristoffersen A, Hausken T. Efficacy of faecal microbiota transplantation for patients with irritable bowel syndrome in a randomised, double-blind, placebo-controlled study. Gut. 2020;69(5):859–67. Epub 2019/12/20. doi: 10.1136/gutjnl-2019-319630. PubMed PMID: 31852769; PMCID: PMC7229896.

31. Hamazaki M, Sawada T, Yamamura T, Maeda K, Mizutani Y, Ishikawa E, Furune S, Yamamoto K, Ishikawa T, Kakushima N, Furukawa K, Ohno E, Honda T, Kawashima H, Ishigami M, Nakamura M, Fujishiro M. Fecal microbiota transplantation in the treatment of irritable bowel syndrome: a single-center prospective study in Japan. BMC Gastroenterol. 2022;22(1):342. Epub 2022/07/15. doi: 10.1186/s12876-022-02408-5. PubMed PMID: 35836115; PMCID: PMC9284895.

32. Holvoet T, Joossens M, Vázquez-Castellanos JF, Christiaens E, Heyerick L, Boelens J, Verhasselt B, van Vlierberghe H, De Vos M, Raes J, De Looze D. Fecal Microbiota Transplantation Reduces Symptoms in Some Patients With Irritable Bowel Syndrome With Predominant Abdominal Bloating: Short- and Long-term Results From a Placebo-Controlled Randomized Trial. Gastroenterology. 2021;160(1):145–57.e8. Epub 2020/07/19. doi: 10.1053/j.gastro.2020.07.013. PubMed PMID: 32681922.

33. Blaser MJ. Fecal Microbiota Transplantation for Dysbiosis - Predictable Risks. N Engl J Med. 2019;381(21):2064–6. Epub 2019/10/31. doi: 10.1056/NEJMe1913807. PubMed PMID: 31665573.

34. DeFilipp Z, Bloom PP, Torres Soto M, Mansour MK, Sater MRA, Huntley MH, Turbett S, Chung RT, Chen YB, Hohmann EL. Drug-Resistant E. coli Bacteremia Transmitted by Fecal Microbiota Transplant. N Engl J Med. 2019;381(21):2043–50. Epub 2019/10/31. doi: 10.1056/NEJMoa1910437. PubMed PMID: 31665575.

35. Allegretti JR, Kassam Z, Mullish BH, Chiang A, Carrellas M, Hurtado J, Marchesi JR, McDonald JAK, Pechlivanis A, Barker GF, Miguéns Blanco J, Garcia-Perez I, Wong WF, Gerardin Y, Silverstein M, Kennedy K, Thompson C. Effects of Fecal Microbiota Transplantation With Oral Capsules in Obese Patients. Clin Gastroenterol Hepatol. 2020;18(4):855–63.e2. Epub 2019/07/14. doi: 10.1016/j.cgh.2019.07.006. PubMed PMID: 31301451.

36. Allegretti JR, Kassam Z, Hurtado J, Marchesi JR, Mullish BH, Chiang A, Thompson CC, Cummings BP. Impact of fecal microbiota transplantation with capsules on the prevention of metabolic syndrome among patients with obesity. Hormones (Athens). 2021;20(1):209–11. Epub 2021/01/10. doi: 10.1007/s42000-020-00265-z. PubMed PMID: 33420959; PMCID: PMC8432937.

37. Mocanu V, Zhang Z, Deehan EC, Kao DH, Hotte N, Karmali S, Birch DW, Samarasinghe KK, Walter J, Madsen KL. Fecal microbial transplantation and fiber supplementation in patients with severe obesity and metabolic syndrome: a randomized double-blind, placebo-controlled phase 2 trial. Nat Med. 2021;27(7):1272–9. Epub 2021/07/07. doi: 10.1038/s41591-021-01399-2. PubMed PMID: 34226737.

38. de Groot P, Scheithauer T, Bakker GJ, Prodan A, Levin E, Khan MT, Herrema H, Ackermans M, Serlie MJM, de Brauw M, Levels JHM, Sales A, Gerdes VE, Ståhlman M, Schimmel AWM, Dallinga-Thie G, Bergman JJ, Holleman F, Hoekstra JBL, Groen A, Bäckhed F, Nieuwdorp M. Donor metabolic characteristics drive effects of faecal microbiota transplantation on recipient insulin sensitivity, energy expenditure and intestinal transit time. Gut. 2020;69(3):502–12. Epub 2019/05/31. doi: 10.1136/gutjnl-2019-318320. PubMed PMID: 31147381; PMCID: PMC7034343.

39. Yu EW, Gao L, Stastka P, Cheney MC, Mahabamunuge J, Torres Soto M, Ford CB, Bryant JA, Henn MR, Hohmann EL. Fecal microbiota transplantation for the improvement of metabolism in obesity: The FMT-TRIM double-blind placebo-controlled pilot trial. PLoS Med. 2020;17(3):e1003051. Epub 2020/03/10. doi: 10.1371/journal.pmed.1003051. PubMed PMID: 32150549; PMCID: PMC7062239 following competing interests: EWY has received a research grant from Amgen Inc. and from Doris Duke Charitable Foundation, outside the submitted work. ELH has served as a consultant to Artugen Therapeutics and Matrivax Research and Development Corporation, and has received a research grant from Kaleido, outside the submitted work. CBF, JAB, and MRH are employees of Seres Therapeutics, Inc. All other authors have declared that no competing interests exist.

40. Vrieze A, Van Nood E, Holleman F, Salojärvi J, Kootte RS, Bartelsman JF, Dallinga-Thie GM, Ackermans MT, Serlie MJ, Oozeer R, Derrien M, Druesne A, Van Hylckama Vlieg JE, Bloks VW, Groen AK, Heilig HG, Zoetendal EG, Stroes ES, de Vos WM, Hoekstra JB, Nieuwdorp M. Transfer of intestinal microbiota from lean donors increases insulin sensitivity in individuals with metabolic syndrome. Gastroenterology. 2012;143(4):913–6.e7. Epub 2012/06/26. doi: 10.1053/j.gastro.2012.06.031. PubMed PMID: 22728514.

41. Gao A, Su J, Liu R, Zhao S, Li W, Xu X, Li D, Shi J, Gu B, Zhang J, Li Q, Wang X, Zhang Y, Xu Y, Lu J, Ning G, Hong J, Bi Y, Gu W, Wang J, Wang W. Sexual dimorphism in glucose metabolism is shaped by androgen-driven gut microbiome. Nat Commun. 2021;12(1):7080. Epub 2021/12/08. doi: 10.1038/s41467-021-27187-7. PubMed PMID: 34873153; PMCID: PMC8648805.

42. Acharya KD, Gao X, Bless EP, Chen J, Tetel MJ. Estradiol and high fat diet associate with changes in gut microbiota in female ob/ob mice. Sci Rep. 2019;9(1):20192. Epub 2019/12/29. doi: 10.1038/s41598-019-56723-1. PubMed PMID: 31882890; PMCID: PMC6934844.

43. Liu H, Shiver AL, Price MN, Carlson HK, Trotter VV, Chen Y, Escalante V, Ray J, Hern KE, Petzold CJ, Turnbaugh PJ, Huang KC, Arkin AP, Deutschbauer AM. Functional genetics of human gut commensal Bacteroides thetaiotaomicron reveals metabolic requirements for growth across environments. Cell Rep. 2021;34(9):108789. Epub 2021/03/04. doi: 10.1016/j.celrep.2021.108789. PubMed PMID: 33657378; PMCID: PMC8121099.

44. Kaliannan K, Robertson RC, Murphy K, Stanton C, Kang C, Wang B, Hao L, Bhan AK, Kang JX. Estrogen-mediated gut microbiome alterations influence sexual dimorphism in metabolic syndrome in mice. Microbiome. 2018;6(1):205. Epub 2018/11/15. doi: 10.1186/s40168-018-0587-0. PubMed PMID: 30424806; PMCID: PMC6234624.

45. Liu R, Hong J, Xu X, Feng Q, Zhang D, Gu Y, Shi J, Zhao S, Liu W, Wang X, Xia H, Liu Z, Cui B, Liang P, Xi L, Jin J, Ying X, Wang X, Zhao X, Li W, Jia H, Lan Z, Li F, Wang R, Sun Y, Yang M, Shen Y, Jie Z, Li J, Chen X, Zhong H, Xie H, Zhang Y, Gu W, Deng X, Shen B, Xu X, Yang H, Xu G, Bi Y, Lai S, Wang J, Qi L, Madsen L, Wang J, Ning G, Kristiansen K, Wang W. Gut microbiome and serum metabolome alterations in obesity and after weight-loss intervention. Nat Med. 2017;23(7):859–68. Epub 2017/06/20. doi: 10.1038/nm.4358. PubMed PMID: 28628112.

46. Takeuchi T, Kubota T, Nakanishi Y, Tsugawa H, Suda W, Kwon AT, Yazaki J, Ikeda K, Nemoto S, Mochizuki Y, Kitami T, Yugi K, Mizuno Y, Yamamichi N, Yamazaki T, Takamoto I, Kubota N, Kadowaki T, Arner E, Carninci P, Ohara O, Arita M, Hattori M, Koyasu S, Ohno H. Gut microbial carbohydrate metabolism contributes to insulin resistance. Nature. 2023;621(7978):389–95. Epub 2023/08/31. doi: 10.1038/s41586-023-06466-x. PubMed PMID: 37648852; PMCID: PMC10499599 metabolic effects of gut bacteria identified by a human cohort. The other authors declare no competing interests.

47. Barnett JA, Gibson DL. H(2)Oh No! The importance of reporting your water source in your in vivo microbiome studies. Gut Microbes. 2019;10(3):261–9. Epub 2018/11/18. doi: 10.1080/19490976.2018.1539599. PubMed PMID: 30442070; PMCID: PMC6546325.

48. Parker KD, Albeke SE, Gigley JP, Goldstein AM, Ward NL. Microbiome Composition in Both Wild-Type and Disease Model Mice Is Heavily Influenced by Mouse Facility. Front Microbiol. 2018;9:1598. Epub 2018/08/07. doi: 10.3389/fmicb.2018.01598. PubMed PMID: 30079054; PMCID: PMC6062620.

49. Karastergiou K, Smith SR, Greenberg AS, Fried SK. Sex differences in human adipose tissues - the biology of pear shape. Biol Sex Differ. 2012;3(1):13. Epub 2012/06/02. doi: 10.1186/2042-6410-3-13. PubMed PMID: 22651247; PMCID: PMC3411490.

50. Perraudeau F, McMurdie P, Bullard J, Cheng A, Cutcliffe C, Deo A, Eid J, Gines J, Iyer M, Justice N, Loo WT, Nemchek M, Schicklberger M, Souza M, Stoneburner B, Tyagi S, Kolterman O. Improvements to postprandial glucose control in subjects with type 2 diabetes: a multicenter, double blind, randomized placebo-controlled trial of a novel probiotic formulation. BMJ Open Diabetes Res Care. 2020;8(1). Epub 2020/07/18. doi: 10.1136/bmjdrc-2020-001319. PubMed PMID: 32675291; PMCID: PMC7368581.

51. Depommier C, Everard A, Druart C, Plovier H, Van Hul M, Vieira-Silva S, Falony G, Raes J, Maiter D, Delzenne NM, de Barsy M, Loumaye A, Hermans MP, Thissen JP, de Vos WM, Cani PD. Supplementation with Akkermansia muciniphila in overweight and obese human volunteers: a proof-of-concept exploratory study. Nat Med. 2019;25(7):1096–103. Epub 2019/07/03. doi: 10.1038/s41591-019-0495-2. PubMed PMID: 31263284; PMCID: PMC6699990.

52. Plovier H, Everard A, Druart C, Depommier C, Van Hul M, Geurts L, Chilloux J, Ottman N, Duparc T, Lichtenstein L, Myridakis A, Delzenne NM, Klievink J, Bhattacharjee A, van der Ark KC, Aalvink S, Martinez LO, Dumas ME, Maiter D, Loumaye A, Hermans MP, Thissen JP, Belzer C, de Vos WM, Cani PD. A purified membrane protein from Akkermansia muciniphila or the pasteurized bacterium improves metabolism in obese and diabetic mice. Nat Med. 2017;23(1):107–13. Epub 2016/11/29. doi: 10.1038/nm.4236. PubMed PMID: 27892954.

53. Paone P, Suriano F, Jian C, Korpela K, Delzenne NM, Van Hul M, Salonen A, Cani PD. Prebiotic oligofructose protects against high-fat diet-induced obesity by changing the gut microbiota, intestinal mucus production, glycosylation and secretion. Gut Microbes. 2022;14(1):2152307. Epub 2022/12/01. doi: 10.1080/19490976.2022.2152307. PubMed PMID: 36448728; PMCID: PMC9715274.

54. Wei X, Yang Z, Rey FE, Ridaura VK, Davidson NO, Gordon JI, Semenkovich CF. Fatty acid synthase modulates intestinal barrier function through palmitoylation of mucin 2. Cell Host Microbe. 2012;11(2):140–52. Epub 2012/02/22. doi: 10.1016/j.chom.2011.12.006. PubMed PMID: 22341463; PMCID: PMC3285413.

55. Wrzosek L, Miquel S, Noordine ML, Bouet S, Joncquel Chevalier-Curt M, Robert V, Philippe C, Bridonneau C, Cherbuy C, Robbe-Masselot C, Langella P, Thomas M. Bacteroides thetaiotaomicron and Faecalibacterium prausnitzii influence the production of mucus glycans and the development of goblet cells in the colonic epithelium of a gnotobiotic model rodent. BMC Biol. 2013;11:61. Epub 2013/05/23. doi: 10.1186/1741-7007-11-61. PubMed PMID: 23692866; PMCID: PMC3673873.

56. Kühn F, Adiliaghdam F, Cavallaro PM, Hamarneh SR, Tsurumi A, Hoda RS, Munoz AR, Dhole Y, Ramirez JM, Liu E, Vasan R, Liu Y, Samarbafzadeh E, Nunez RA, Farber MZ, Chopra V, Malo MS, Rahme LG, Hodin RA. Intestinal alkaline phosphatase targets the gut barrier to prevent aging. JCI Insight. 2020;5(6). Epub 2020/03/28. doi: 10.1172/jci.insight.134049. PubMed PMID: 32213701; PMCID: PMC7213802.

57. Bates JM, Akerlund J, Mittge E, Guillemin K. Intestinal alkaline phosphatase detoxifies lipopolysaccharide and prevents inflammation in zebrafish in response to the gut microbiota. Cell Host Microbe. 2007;2(6):371–82. Epub 2007/12/15. doi: 10.1016/j.chom.2007.10.010. PubMed PMID: 18078689; PMCID: PMC2730374.

58. Malo MS. A High Level of Intestinal Alkaline Phosphatase Is Protective Against Type 2 Diabetes Mellitus Irrespective of Obesity. EBioMedicine. 2015;2(12):2016–23. Epub 2016/02/05. doi: 10.1016/j.ebiom.2015.11.027. PubMed PMID: 26844282; PMCID: PMC4703762.

59. Smith PM, Howitt MR, Panikov N, Michaud M, Gallini CA, Bohlooly YM, Glickman JN, Garrett WS. The microbial metabolites, short-chain fatty acids, regulate colonic Treg cell homeostasis. Science. 2013;341(6145):569–73. Epub 2013/07/06. doi: 10.1126/science.1241165. PubMed PMID: 23828891; PMCID: PMC3807819.

60. Arpaia N, Campbell C, Fan X, Dikiy S, van der Veeken J, deRoos P, Liu H, Cross JR, Pfeffer K, Coffer PJ, Rudensky AY. Metabolites produced by commensal bacteria promote peripheral regulatory T-cell generation. Nature. 2013;504(7480):451–5. Epub 2013/11/15. doi: 10.1038/nature12726. PubMed PMID: 24226773; PMCID: PMC3869884.

61. Komano H, Fujiura Y, Kawaguchi M, Matsumoto S, Hashimoto Y, Obana S, Mombaerts P, Tonegawa S, Yamamoto H, Itohara S, et al. Homeostatic regulation of intestinal epithelia by intraepithelial gamma delta T cells. Proc Natl Acad Sci U S A. 1995;92(13):6147–51. Epub 1995/06/20. doi: 10.1073/pnas.92.13.6147. PubMed PMID: 7597094; PMCID: PMC41659.

62. Boismenu R, Havran WL. Modulation of epithelial cell growth by intraepithelial gamma delta T cells. Science. 1994;266(5188):1253–5. Epub 1994/11/18. doi: 10.1126/science.7973709. PubMed PMID: 7973709.

63. Arora S, Thompson PJ, Wang Y, Bhattacharyya A, Apostolopoulou H, Hatano R, Naikawadi RP, Shah A, Wolters PJ, Koliwad S, Bhattacharya M, Bhushan A. Invariant Natural Killer T cells coordinate removal of senescent cells. Med. 2021;2(8):938–50. Epub 2021/10/08. doi: 10.1016/j.medj.2021.04.014. PubMed PMID: 34617070; PMCID: PMC8491998.

64. Mousa A, Naderpoor N, Mellett N, Wilson K, Plebanski M, Meikle PJ, de Courten B. Lipidomic profiling reveals early-stage metabolic dysfunction in overweight or obese humans. Biochim Biophys Acta Mol Cell Biol Lipids. 2019;1864(3):335–43. Epub 2018/12/27. doi: 10.1016/j.bbalip.2018.12.014. PubMed PMID: 30586632.

65. Mendham AE, Goedecke JH, Zeng Y, Larsen S, George C, Hauksson J, Fortuin-de Smidt MC, Chibalin AV, Olsson T, Chorell E. Exercise training improves mitochondrial respiration and is associated with an altered intramuscular phospholipid signature in women with obesity. Diabetologia. 2021;64(7):1642–59. Epub 2021/03/27. doi: 10.1007/s00125-021-05430-6. PubMed PMID: 33770195; PMCID: PMC8187207.

