## Supplemental Figures for "Beneficial metabolic effects of PAHSAs depend on the gut microbiota in diet-induced obese mice"

**Figure S1, related to Figure 1. PAHSA treatment in mice increases insulin sensitivity and alters the gut microbiota and plasma and cecal metabolomes.**

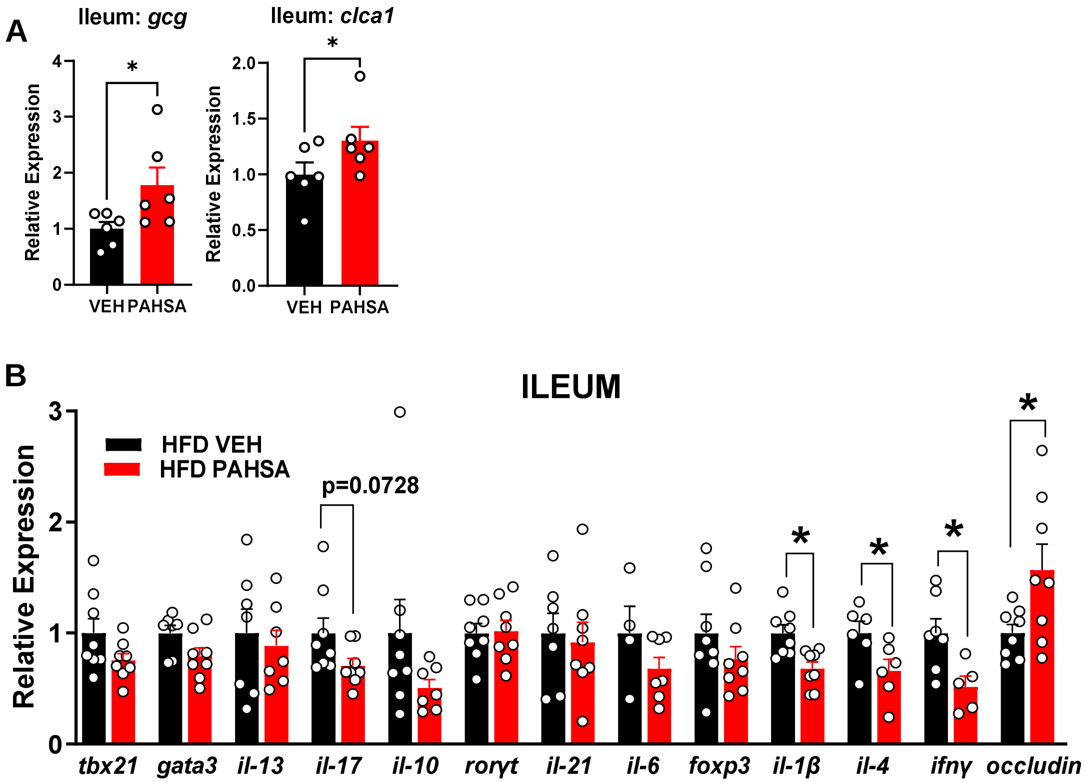

**Figure S2, related to Figure 2. The beneficial PAHSAs effects on glucose homeostasis are transmissible by fecal microbiota transplantation.**

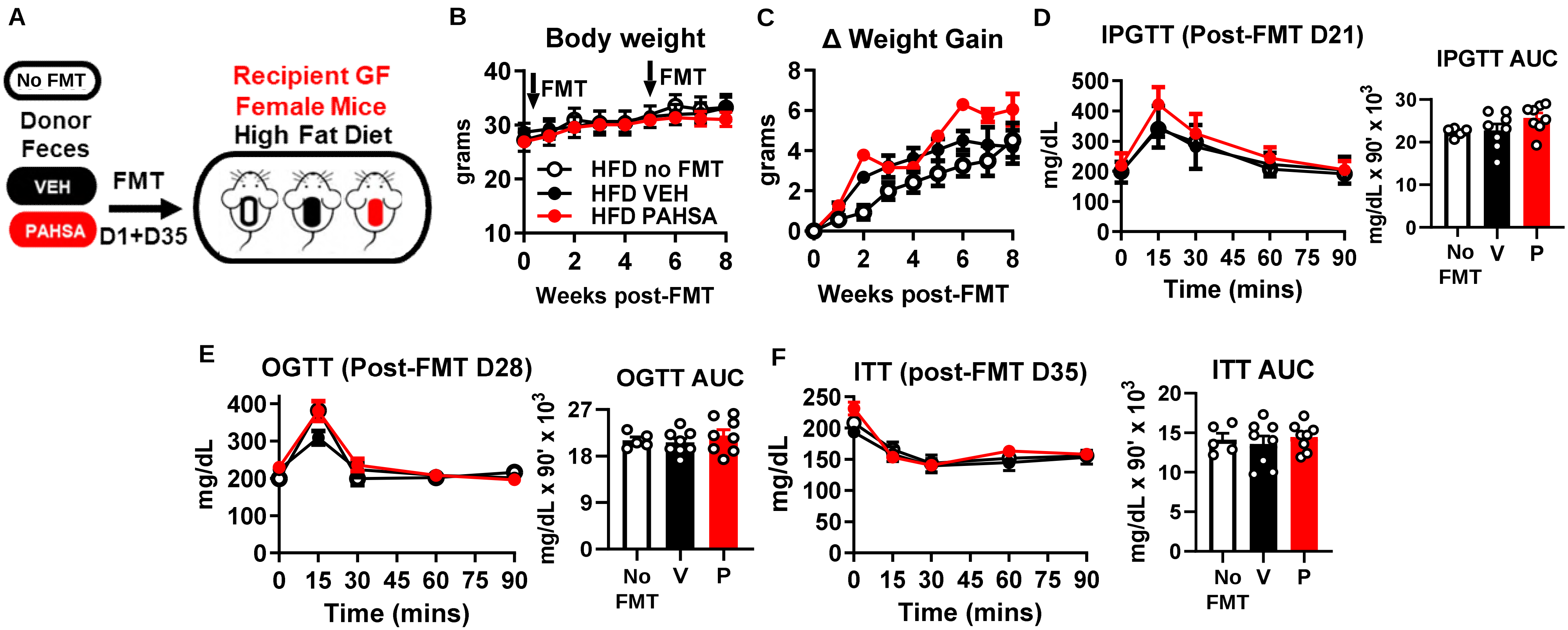

**Figure S3, related to Figure 3. The gut microbiota is necessary for PAHSAs to improve glucose homeostasis in HFD-fed mice.**

**A**

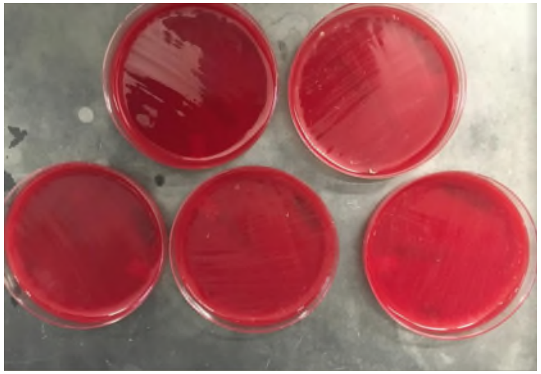

**Bacterial cultures of mouse feces collected from HFD Gnotobiotic isolators**

**Figure S4, related to Figure 4. The gut microbiome has sex-specific responses to *Bacteroides thetaiotaomicron* supplementation in dietary obese mice.**

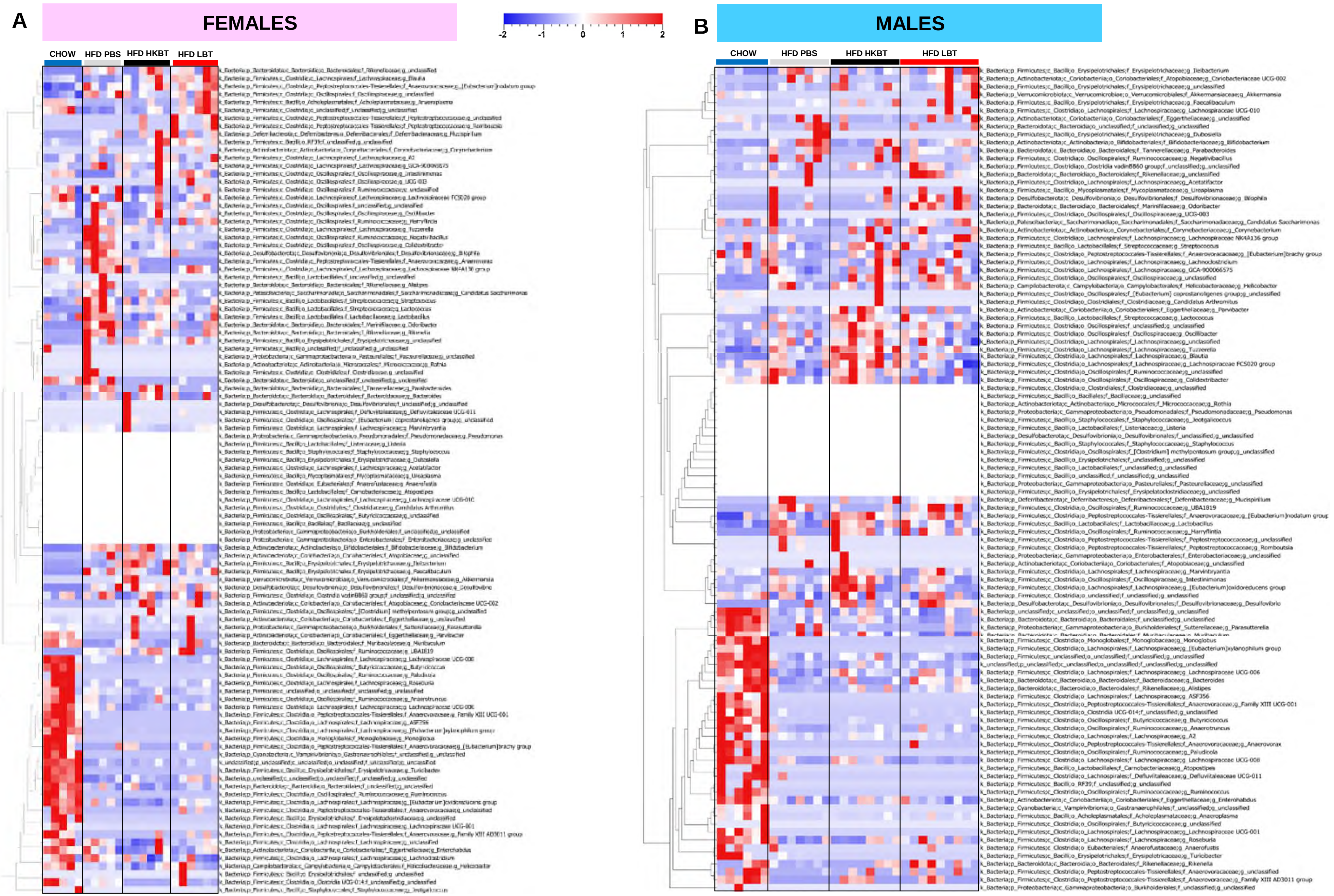

**Figure S5, related to Figure 5. *Bacteroides thetaiotaomicron* supplementation improves host metabolism in dietary obese mice.**

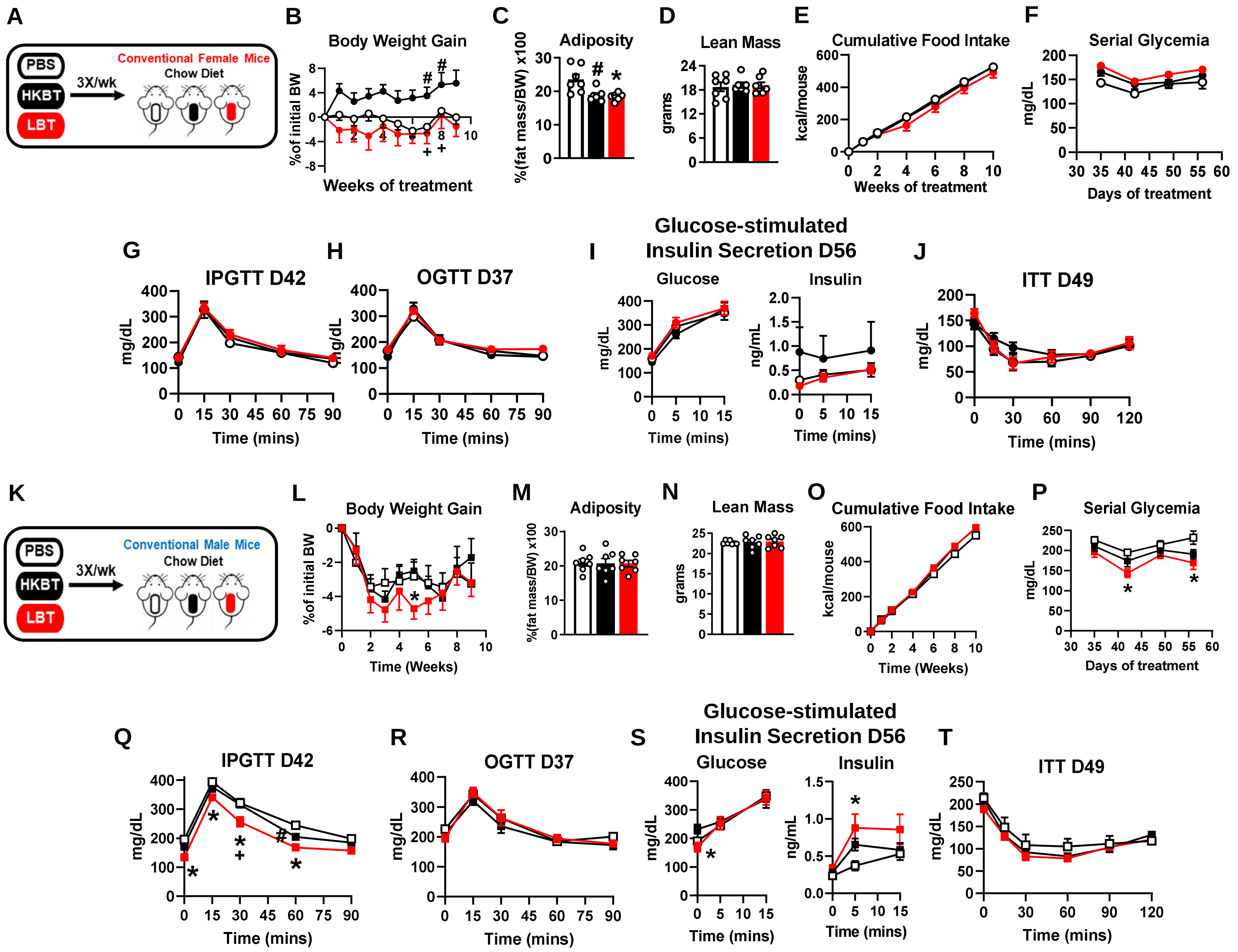

**Continued Figure S5, related to Figure 5. *Bacteroides thetaiotaomicron* supplementation improves host metabolism in dietary obese mice.**

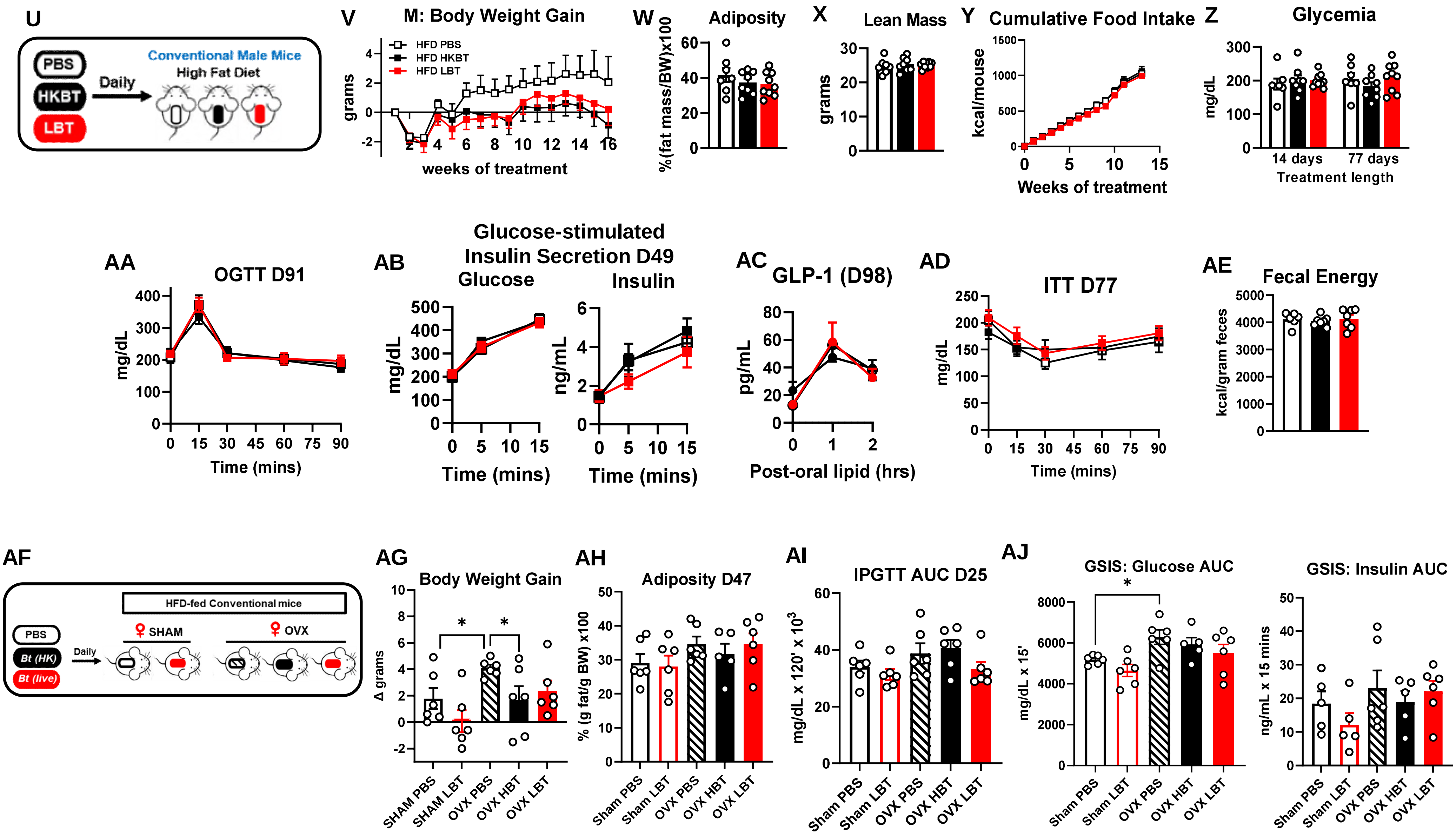

**Figure S6, related to Figure 6. Effects of *Bt* supplementation on the gut mucosa and intestinal innate immune cells in dietary obese mice**

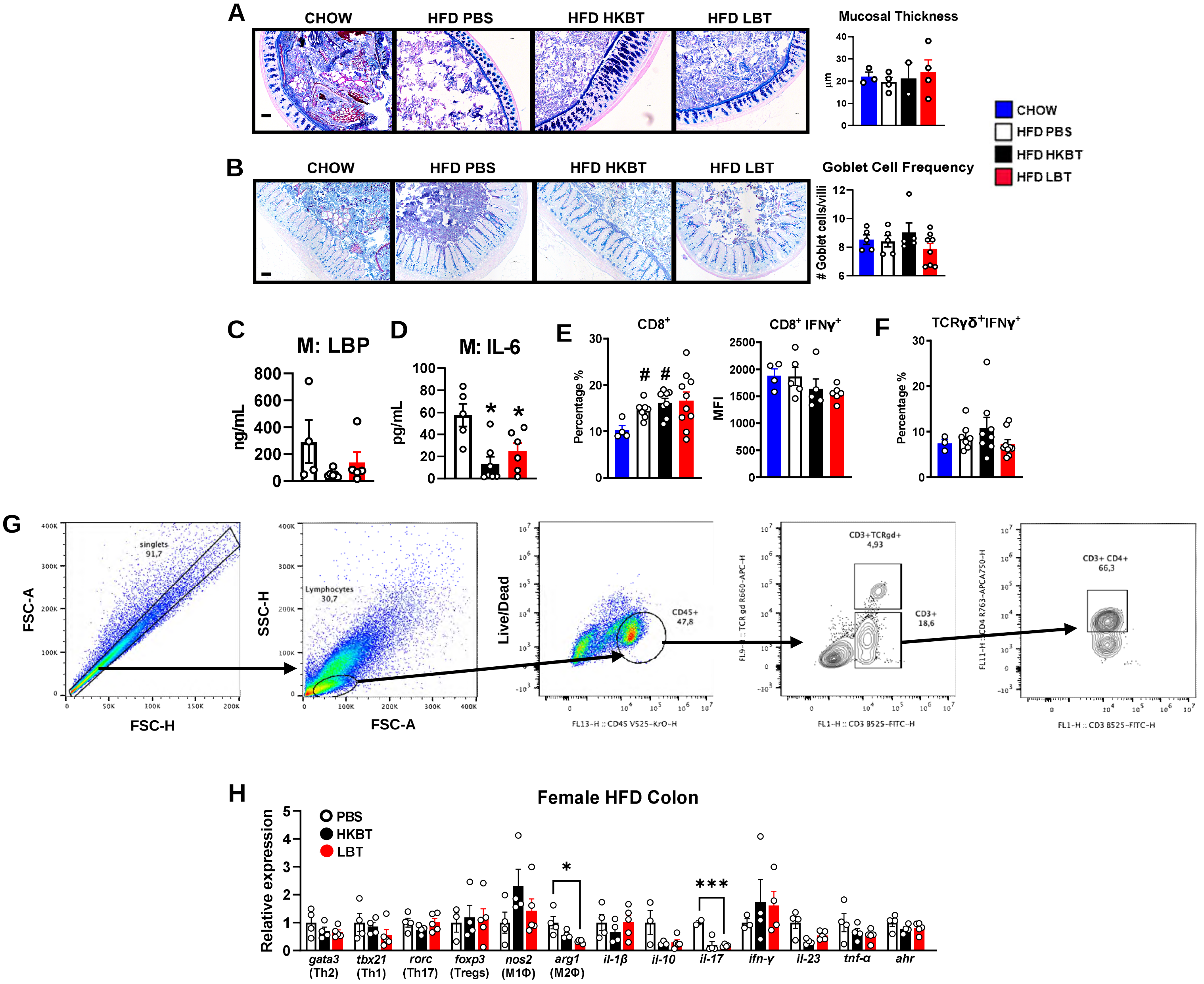
